# A multi-layered systems approach for renal cell carcinoma

**DOI:** 10.1101/2020.01.13.904235

**Authors:** Lindsay S. Cooley, Justine Rudewicz, Wilfried Souleyreau, Kim Clarke, Francesco Falciani, Maeva Dufies, Stephanie Verbeke, Andrea Emanuelli, Sebastien Benzekry, Cécile Taing, Emeline Julie Ribot, Sylvain Miraux, Diether Lambrechts, Elodie Modave, Raphael Pineau, Marie-Alix Derieppe, Damien Ambrosetti, Jean-Christophe Bernhard, Alain Ravaud, Sylvie Négrier, Jean-Marc Ferrero, Gilles Pagès, Macha Nikolski, Andreas Bikfalvi

**Author notes:** Correspondence: Andreas Bikfalvi MD PhD, LAMC INSERM U1029, Université Bordeaux Allée Geoffroy St Hilaire 33615 Pessac France. These authors have equally contributed to this work (Lindsay S. Cooley, Justine Rudewicz, Wilfried Souleyreau).

## Abstract

Renal cell carcinoma (RCC) still lacks prognostic and predictive biomarkers to monitor the disease and the response to therapy. The usual strategy in translational research is to start from human samples, to identify molecular markers and gene networks and then to functionally validate them *in vitro* and in animal models. We devised herein a completely opposite strategy from “mouse to man” by performing an aggressiveness screen and used functional genomics, imaging, clinical data and computational approaches in order to discover molecular pathways and players in renal cancer development and metastasis. Multiple cell lines for primary tumor growth, survival in the blood circulation and lung metastasis or metastatic spread from the primary tumor were generated and analyzed using a multi-layered approach which includes large-scale transcriptome, genome and methylome analyses. Transcriptome and methylome analyses demonstrated distinct clustering in three different groups. Remarkably, DNA sequencing did not show significant genomic variations in the different groups which indicates absence of clonal selection during the *in vivo* amplification process. Transcriptome analysis revealed distinct signatures of tumor aggressiveness which were validated in patient cohorts. Methylome analysis of full-length DNA allowed clustering of the same groups and revealed clinically relevant signatures. Furthermore, we identified SAA2 and CFB as soluble prognostic and predictive biomarkers of the therapeutic response. We also uncovered IL34 as another soluble prognostic biomarker and key regulator of renal cell carcinoma (RCC) progression. This was also functionally validated *in vivo,* and a mathematical model of IL34-dependent primary tumor growth and metastasis development was provided. These results indicate that such multilayered analysis in a RCC animal model leads to meaningful results that are of translational significance.

**One Sentence Summary:** An aggressiveness screen with multilayer systems analysis to identify signatures and biomarkers for renal cell carcinoma aggressiveness.

## INTRODUCTION

Renal Cell Carcinoma (RCC) encompasses a heterogeneous group of cancers derived from renal tubular epithelial cells, including multiple histological and molecular sub-types, of which clear cell RCC (ccRCC) is the most common *(Hsieh et al, 2017)*. The incidence and prevalence of RCC are rising *(Ljungberg et al, 2011; American Cancer Society. Key Statistics about kidney cancer, 2016)*, along with increases in related risk factors such as hypertension, diabetes and obesity *(Chow et al, 2010)*. Gandaglia et al. *(Gandaglia et al, 2014)* reported a continuing upward trend in both incidence and mortality even in patients with localized disease. When disease is localized to the kidney, surgical resection is the preferred option and the prognosis is favorable. However, therapeutic options for metastatic disease are limited. ccRCC metastasizes primarily to the lungs (secondarily to liver and bone), and 5-year survival ranges 10-12 % *(Tsui et al, 2000; Lam et al, 2005; Antonelli et al, 2007)*. Furthermore, 40% of patients with seemingly localized disease later relapse with localized or metastatic disease. Localized recurrence is also difficult to treat and difficult to predict, and has a poor prognosis *(Ö. & Ö., 2016; Chin et al, 2006)*.

Challenges associated with treatment of RCC include high levels of resistance to traditional chemotherapeutic drugs *(Chow et al, 2010)*. The majority of currently available targeted therapies focus on inhibiting angiogenesis driven by the VEGF/VEGFR axis *(Molina & Motzer, 2011)*. More recently, immunotherapeutic agents (nivolumab and ipilimimab) have also been added as first line treatment *(Motzer et al, 2018; Escudier et al, 2019)*. However, such therapies are rarely curative, and eventual drug resistance is almost inevitable. Furthermore, clinical treatment of RCC is hampered by a lack of relevant biomarkers. Currently, no fully validated molecular biomarkers for RCC are used in clinical practice. Response to available treatments and long-term disease-free survival is highly variable and unpredictable. Patient diagnosis, prognosis and clinical decisions are currently (based on histological information such as Fuhrman grade and tumor stage, and are included in the SSIGN or UISS classification for primary tumor risk of recurrence *(Zisman et al, 2002; Patard et al, 2004; Ficarra et al, 2006; Zigeuner et al, 2010)*, and in the MSKCC and/or IMDC classification for metastatic disease *(Heng et al, 2014; Bensalah et al, 2016, 2018)*. In addition, therapy selection is based on limited guidelines and response to previous treatments. In this respect, clinical treatment of RCC lags behind other cancers for which molecular knowledge is invaluable in guiding clinical decisions (e.g. hormone receptor status in breast cancer).

Tumor progression from initiation to full metastasis is a multi-step process and occurs via a series of overlapping stages. Tumor progression can be seen as an evolutionary process whereby tumor cells within the primary tumor adapt to varied selection pressures. Cells must detach from the primary tumor, gain access to and survive in the circulation, exit the vasculature, and survive and proliferate in the environment of the secondary organ. Thus, different mechanisms come into play at different stages, and overall tumor progression is the sum of these processes. A better understanding of the molecular changes that enable cancer cells to overcome the barriers imposed during the metastatic process could aid in the diagnosis, prevention and treatment of metastatic cancer.

The process of serial implantation of a cell line *in vivo* is a known method to render a cell line increasingly aggressive *(Fidler, 1973; Bruns et al, 1999)*. In this study, we leveraged this technique to generate increasingly aggressive sub-lines of mouse renal carcinoma. We then devised a multi-layer systems strategy to generate signatures able to predict global, disease-free and progression-free survival in human patients. Furthermore, we identified and investigated a number of genes with the potential to be biomarkers and therapeutic targets.

## RESULTS

### Generation and characterization of the mouse model

Our strategy is based on a syngeneic mouse renal carcinoma model. When RENCA *(Murphy & Hrushesky, 1973)*, expressing GFP are implanted orthotopically under the kidney capsule in female BALB/c mice, metastasis occurs specifically in the lungs after 14-16 days, with ethical sacrifice at 26 days following implantation (Fig. 1A). We adapted this model to produce new sub-cell lines with progressively enhanced aggressiveness and metastatic potential with differing characteristics. We serially passaged the cell lines using different implantation strategies designed to replicate different aspects of tumor development and metastasis (Fig. 1A). After each passage, primary and metastatic tumor cells were purified from organ explant cultures. The three injections modes were as follows:

i. Orthotopic injection under the renal capsule leading to formation of a primary tumor (“Kidney” group). The primary tumor was explanted and cell lines purified for re-implantation into the kidney. The “Kidney” cell lines thus derived have undergone repeated cycles of primary tumor formation..
ii. Intravenous injection (“Tail vein” group). Tumor cells were injected directly into the blood stream leading to formation of tumors in the lungs in the absence of a tumor in the kidney. The “Tail” cell lines thus derived have undergone repeated cycles of survival in the blood stream, evasion of host immune response, colonization and growth in the lungs.
iii. Orthotopic injection under the renal capsule followed by metastasis to the lungs (“Lung” group). Tumor cells were purified from the lung metastases and re-implanted under the kidney capsule for the subsequent passage. These “Lung” cell lines have undergone the full cycle of primary tumor formation, dissemination in the blood, and secondary tumor formation at a distant site.

**Fig. 1.**
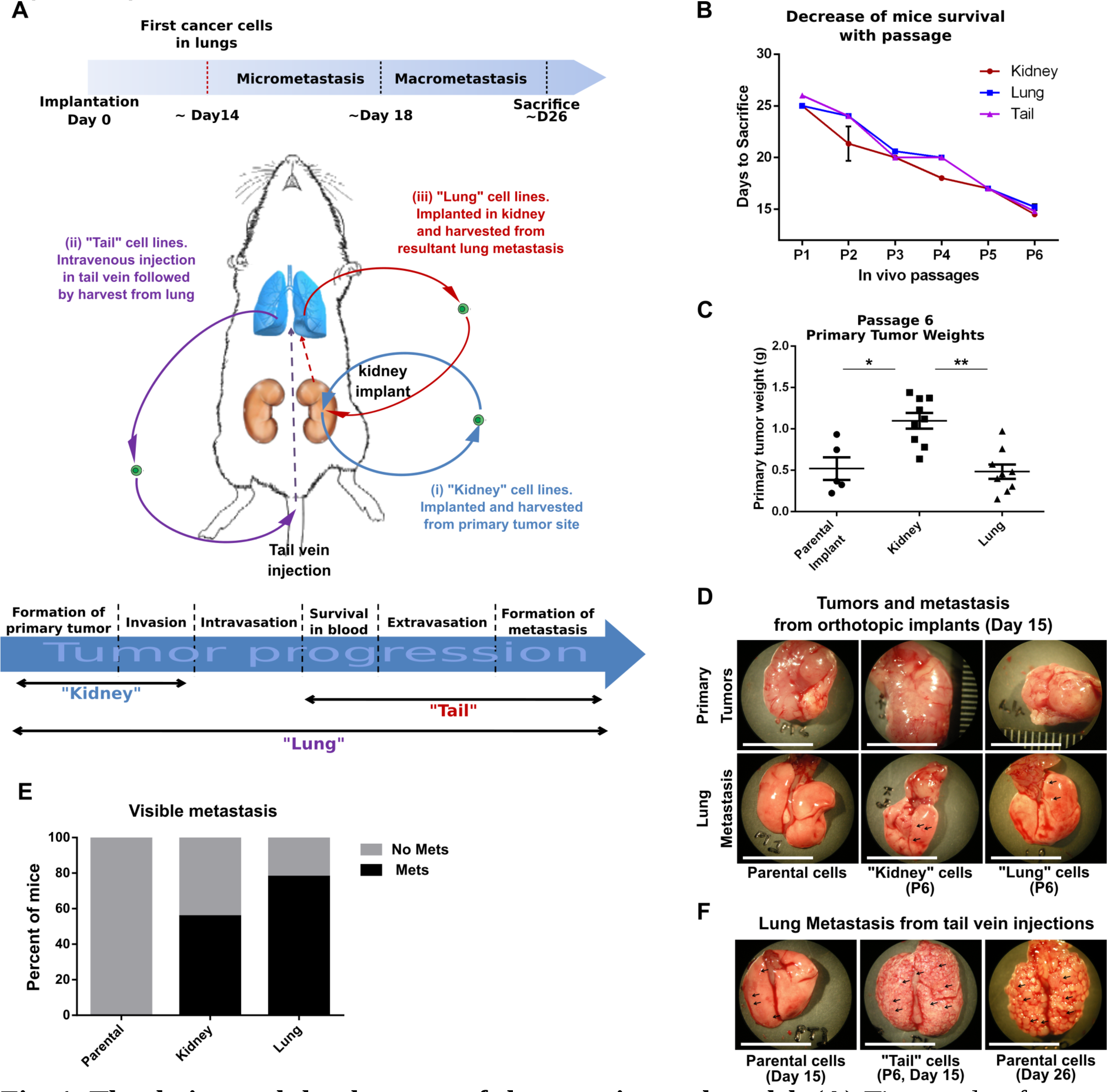
The design and development of the experimental model. (**A**) Time scale of tumor metastasis in mice implanted orthotopically with RENCA cells. Three different injection modalities were used to develop increasingly aggressive cell lines: Kidney (i), expected to reveal enhancement of mechanisms related to primary tumor formation; Tail (ii), expected to recapitulate key aspects that precede metastasis formation (i.e. survival in the blood stream, evasion of host immune response, and distant organ colonization); Lung (iii), representing cells which underwent the full cycle of primary tumor formation, dissemination, and secondary tumor formation at a distant site. At the bottom, scheme of different tumor progression phases and mechanisms which are expected to be enhanced in the relevant cell line. (**B**) the graph shows reduced mice survival, represented as time passed since the implantation until the day of sacrifice, after 6 cycles of RENCA injection. Data are represented as mean ± SEM. (**C**) Dots graph shows the weight of primary tumors generated by parental (P0), or late passaged Kidney P6 or Lung P6 RENCA cells, after implantation into kidney. Data are represented as mean ± SEM. Stats ANOVA followed with Tukey’s multiple comparisons. *p<0.05; **p<0.01. (**D**) Representative pictures of primary tumors and lung metastasis generated by orthotopic implant into kidney of either parental (P0) or late passaged cell lines (Kidney P6 or Lung P6). Bar = 1cm. (**E**) Bars graph showing the percentage of mice presenting visible metastases in the lungs, upon injection of late passaged RENCA (either Kidney P6 or Lung P6) compared to parental (P0) cell line. (**F**) representative pictures of lung metastases formed, at day 15, by late passaged RENCA cells (Tail P6), compared to parental cells (P0), which were tail injected. To note, upon tail injection, parental cell line P0 requires a wider window of time (i.e. 26 days) to form visible lung metastases.

After 6 sequentially *in vivo* passages using multiple mice per injection mode and per passage, 67 newly derived sub-cell lines were obtained. After each passage a reduced survival time for the mice of all groups was observed indicating increased agressiveness (Fig. 1B). At day 15 post-implantation, we observed that (i) at passage 6, “Kidney” cell lines generated significantly larger primary tumors than parental or “Lung” cell lines (Fig. 1, C and D); (ii) visible metastases were detected in the lungs of “Kidney” and of “Lung” cell implanted mice (Fig. 1, D and E), whereas lungs of mice with orthotopic implant of parental cells showed no macroscopic metastases indicating that metastatic ability was enhanced in passaged cell lines. ***(i)*** (iii) in the intravenous injection experimental metastasis model (i.e. “Tail” cell line), the number and size of metastases were drastically increased (Fig. 1F) compared to the parental cells, as a consequence of enhanced aggressiveness and specialization of serial passaged cells Thus, serial passaging resulted in generation of cell lines that were more aggressive and specialized when compared to control cells.

### Functional genomics of amplified mouse cell lines

#### Transcriptomic analysis

We performed full genome transcriptomic analysis of the 67 cell lines. The P1 cell line was excluded from the analysis because of insufficient number of animals (<3). Therefore, data acquired from the different lines were labeled S0 to S5 (S0 being the parental cells and S1 to S5 representing P2 to P6). We used Principal Component Analysis (PCA) in order to summarize the information contained in our data sets for sample series S2 and S5. Principal Component Analyses (showed that, while the subtypes had similar expression profiles in the early passages, the transcriptional profiles became different between “Kidney” (K), “Tail” (T), and “Lung” (L) groups in later passages and, thus, clustered into distinct groups (Fig. 2, A and B). Tight clustering of the biological replicates (one mouse per data point) also showed that the profiles were stable. When all genes that were major contributors to the Principal Component 1 (PC1) and 2 (PC2) were pooled, the heat map of the transcriptomic profiles revealed a gradual change in expression (Fig. 2C).

**Fig. 2.**
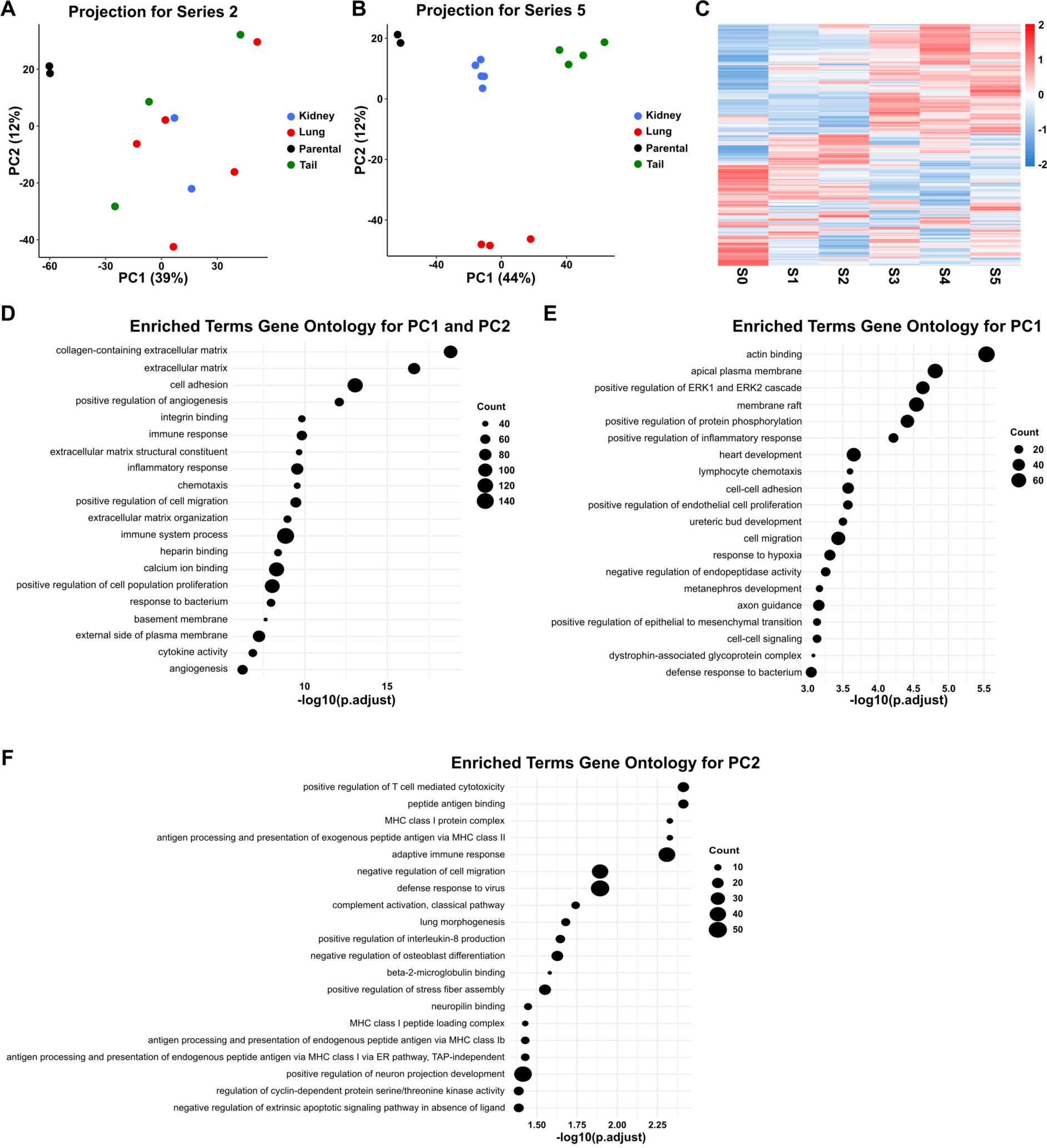
Analysis of transcription data sets. (**A** and **B**) Projection of samples onto principal component (PC) 1 and 2 for series 2 and 5 respectively. Samples of series 2 did not cluster whereas series 5 did. **c** Heatmap for PC1-et PC2-associated gene expression from the PCA analysis through S0 to S5. A progressive pattern is displayed. (**D** to **F**) Each enriched GO term is represented in function of the -log10 of its adjusted p-value. Count: number of genes in the GO term. Intersection of enriched GO terms for PC1-andPC2-associated genes from the PCA analysis for S5 (**D**); Unique enriched GO terms for PC1 (**E**) or PC2 (**F**) associated genes from PCA analysis for S5.

Enrichment analysis of these most contributing genes (see Methods for details) demonstrated several highly enriched categories. These include the following GO terms: extracellular matrix, angiogenesis, cell proliferation, cell adhesion, cell migration, immune process and inflammation and apoptosis. Fig. 2, D to F represent enriched GO terms in common for PC1 and PC2 or specific to PC1 or PC2, respectively.

Next, we investigated whether transcriptional signatures derived from the differentially expressed genes in the K, T and L groups could predict outcome for patients using the Clear Cell Renal Cell Carcinoma dataset (KIRC) from The Cancer Genome Atlas (TCGA).

The general strategy is outlined in fig. S1. We compared genes that changed their expression between the parental and S5 cell line and named these genes as progressively regulated (up or down) (table S1). We included in our analysis only genes having their expression consistently increasing in the different series.

**Table 1.**
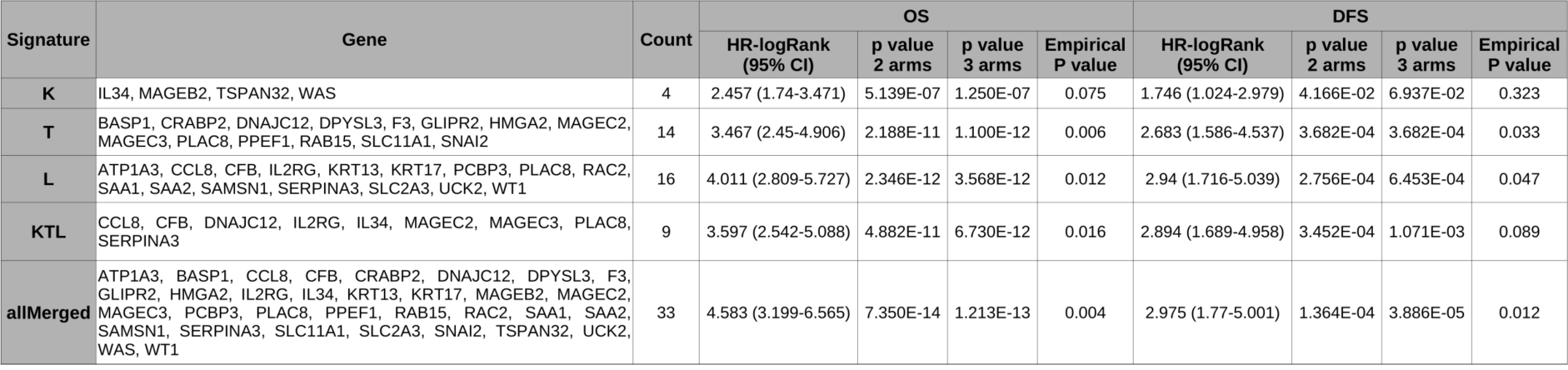
Summary table of the signatures and their predictive value in the KIRC TCGA cohort.

The analysis has been performed by comparison to the parental cell line for K, T and L groups alone. The additional group “Kidney-Tail-Lung” (KTL) is represented by pooling the different groups in a unique group. The KTL group is also compared to the parental cell line (see Methods for details). The table S1 lists genes differentially expressed in each group (K, T, L, KTL).

A total of 131 genes for all groups satisfied the inclusion criteria. Among these, 26 were in the K, 45 in the T, 59 in the L and 31 in the KTL group (table S1). Among those, 123 genes were upregulated and only a small fraction was downregulated.

We next investigated the predictive value of these 4 gene signatures using TCGA (KIRC cohort). Since only 8 genes were downregulated, we chose only to include the upregulated genes in the validation of our signatures. Table S2 depicts the list of genes for each group. For each gene, we fitted a Cox-proportional hazard regression model based on overall survival (OS) or disease-free survival (DFS). A gene was conserved if its false-discovery rate (FDR) adjusted p-value of its log-rank test was lower than 0.01 and if the hazard ratio was in agreement with the differential expression. We have further created a 5th “All Merged” signature composed of genes that are members of all of the previously defined 4 signatures.

Table 1 depicts the signatures for each group (K, T, L, KTL) and their significance for OS and DFS. In Fig. 3 the results of “Lung” subgroup are depicted. We analyzed OS (Fig. 3A) of all patients and DFS (Fig. 3B) of M0 patients. We validated our signatures by computing an empirical p-value and by testing our signature against 1000 random signatures of equivalent size (Fig. 3,C and D). Furthermore, we performed multivariate Cox regression analysis of our signature (Fig. 3E and F). After adjusting for clinical variables (TNM stage and Fuhrman grade), the Lung signature remained an independent prognostic factor for predicting both OS and DFS. The results for the other signatures are shown in fig. S2 to S5. The Lung signature remains the most significant for OS and DFS when compared to the others, except for the “All Merged” signature. The performances of the signatures were the following for the different groups when ranked according to the hazard ratio: K<T<KTL<L<All Merged for both OS and DFS.

**Fig. 3.**
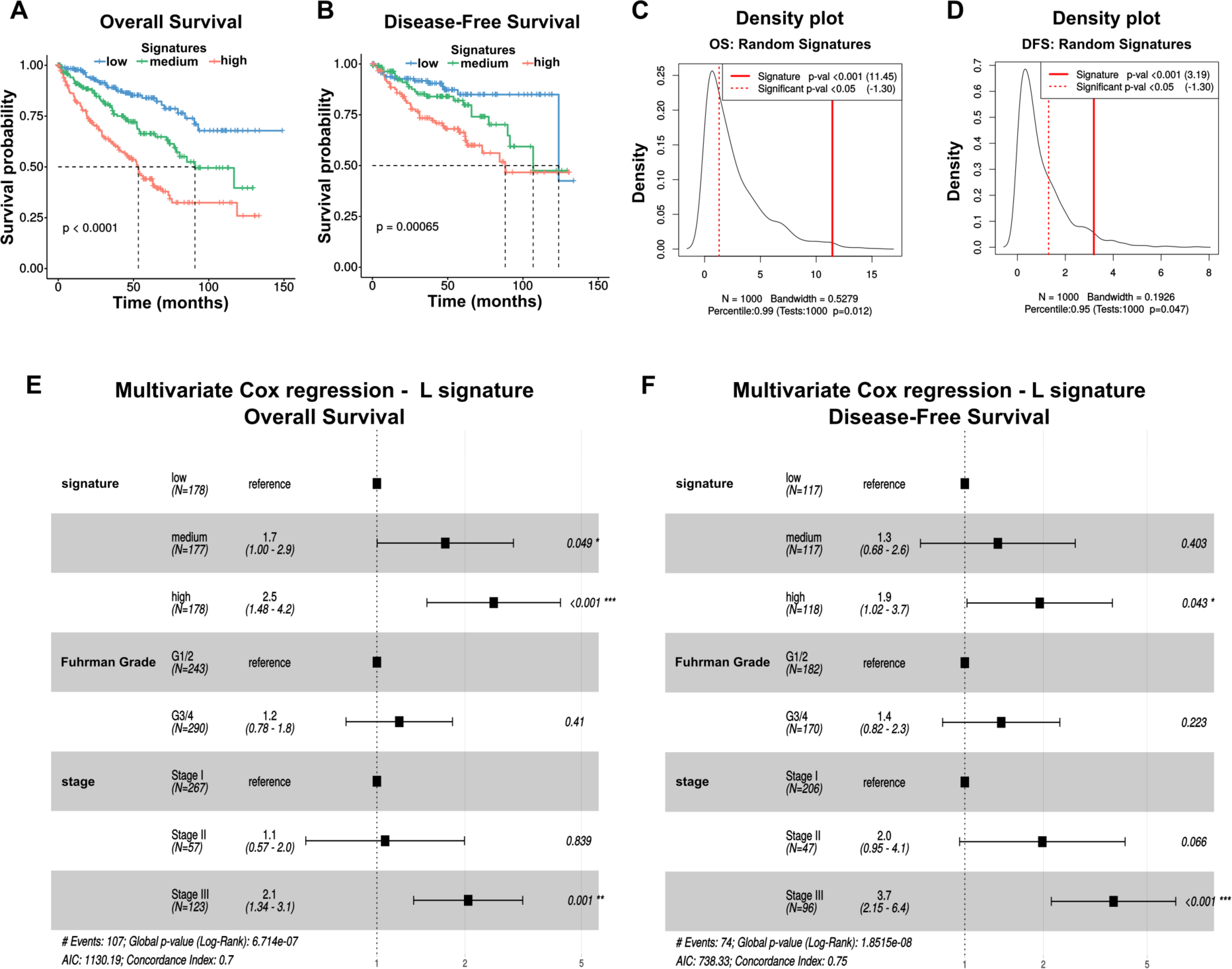
Validation of the “Lung” signature. (**A** and **B**) Kaplan-Meier for overall (OS) and disease-free (DFS) survival analysis stratified in 3 groups of equivalent size. Signature “low”: patient group with low score; signature “medium”: patient group with a medium score; signature “high”: patient group with high score. (**C** and **D**) Density plot of p values (log-rank test) from 1000 random signatures of equal size for OS and DFS. p = empirical p-value. (**E** and **F**) Forest plot of multivariate Cox proportional hazards model for the Lung signature in the KIRC cohort adjusted for clinical variables (Fuhrman grade and TNM stage).

The identification of signatures predictive of patient outcome also validates our experimental approach and shows that the strategy of generation of increasingly specialized mouse cell lines revealed novel genes and signatures with relevance to human RCC.

#### DNA sequencing analysis

We subjected both the parental RCC line and those that were passaged 6 times (series S5) to low-coverage whole-genome sequencing to assess whether copy number variability could possibly underlie the change in phenotype (fig. S6). We failed however to observe significant differences in copy numbers between parental and passaged samples, both at the level of the number of breakpoints detected (45 for parental versus 41.75±6.32 for passaged lines) and the percentage of the genome with a copy number different than 2 (19% for parental versus 18%±1% for passaged lines). Visually, we also failed to observe new copy number events. When using ABSOLUTE to estimate purity (proportion of cancer cells present in the sample) and mean ploïdy of the samples, no differences were noted (purity and ploïdy in parental line were of 63% and 2.69, respectively versus 64%±2% and 2.65±3% for passaged lines). This suggests the altered profile of the “Kidney”, “Tail” and “Lung” lines was not due to clonal selection.

#### Methylome sequencing analysis

Since DNA sequencing did not provide conclusive evidence for clonal selection, we conducted a methylome analysis by full methylome sequencing of the S5 series samples for the K, T and L groups as well as of the parental control cells. Similarly to the transcriptional data, Principal Component Analysis (PCA) showed clustering of the global methylome in 4 clusters corresponding to the K, T, L and parental cell line samples (Fig. 4A). Enrichment analysis demonstrated several highly enriched categories (Fig. 4B). These include cell cytokine activity, nervous system development, regulation of cell population/proliferation, extracellular matrix, transcription and gene regulation, angiogenesis, regulation of gene expression among the most significantly enriched categories.

**Fig. 4.**
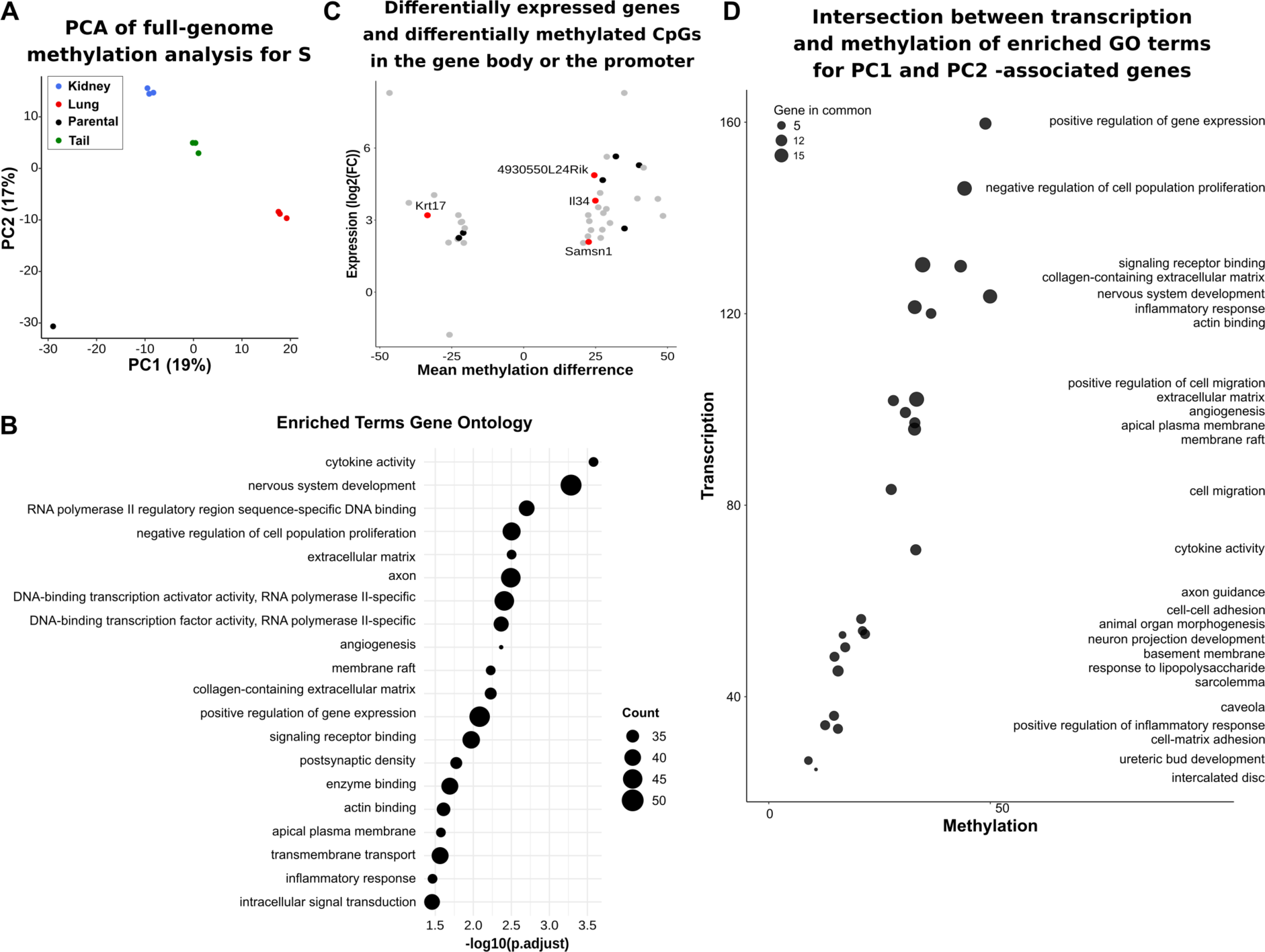
Analysis of methylome data sets. (**A**) Principal component of full-genome methylation analysis for S5 series. (**B**) Enriched GO terms for PC1/PC2-associated genes from the PCA analysis of the S5 methylation series. Each enriched GO term is represented in function of the -log10 of its adjusted p-value. Count: number of genes in the GO term; (**C**) Dot-plot of differentially expressed genes (DEG) corresponding to differentially methylated CpGs in the gene body or the promoter. (**D**) Intersection between transcription and methylation of enriched GO terms for PC1 and PC2-associated genes.

We focused on CpGs that were located in the gene body with methylation equal or higher than 20 % and with a q-value of 0.05. We found 33 095 CpGs differentially methylated, out of which 9 731 were associated with a known gene. They were confronted with the transcriptome results (Fig. 4C). Differentially expressed and methylated genes were included in this analysis if they contained at least one differentially methylated CpG in their gene-body. Several genes contained in the transcriptome signatures were found in this analysis. For progressive signatures (table S1): *Gm2012* (K), *B3galt1* (T), *Krt20* (T), *Thbd* (T), *Adam22* (L), *Apbb1ip* (L), *Arap3* (L), *Wscd2* (L). For final signatures (Table 1 and table S2, corresponding to red dots): *IL34* (KTL), *Rik/MAGEC2/3* (T), *Krt17* (L), *Samsn1* (L).

We then investigated the clinical relevance of CpG methylation in the gene body of these genes in the KIRC TCGA cohort. Two CpGs (cg01782798 and cg26831220) in the gene body of *IL34* were predictive for OS (fig. S7). In our data sets, the differentially methylated cytosine (DMC) was located in intron 1 of *IL34* (fig. S7A), as were cg01782798 and cg26831220 (fig. S7B). According to ENSEMBL for the GRCm38 mouse genome and GeneHancer for the GRCh37 Human genome, these three CpGs are part of enhancer regions. Correlations of *IL34* transcription levels and cg01782798 or cg26831220 are depicted in fig. S7, C and D. Kaplan-Meier and log-rank test for OS were done by separating samples into 2 groups by the median value for each CpG resulting in a p-value of 1.62e-05 and 0.00057 and a Hazard Ratio (HR) of 2.422 (1.651-3.551 95% CI) and 1.970 (1.342-2.894 95% CI) for CpG cg01782798 and cg26831220, respectively (fig. S7, E to G). In the same manner as for the transcriptomics data, the clinical relevance and prognostic performance of the 2 CpGs were tested against 1000 random 1-CpG signatures resulting in empirical p-values of 0.015 and 0.033 for CpG cg01782798 and cg26831220, respectively (fig. 7G).

Since the differential analyses of methylation presents only few similarities with the transcriptomic results, we checked whether these modifications/dis-regulations are related to the same processes. To this aim, we compared enrichment results (GO terms) obtained in the transcriptome and methylome analyses (Fig. 4D). This comparison revealed that in both data sets, a limited number of GO terms are shared (26 GO terms in common of a total of 160 for the transcriptome and 112 for the methylome). This suggests that gene expression and methylation analysis converge only partially to a common set of processes (26/160 = 16% and 26/112=23%).

### Novel RCC biomarkers selected from the signatures and clinical relevance

To identify novel potential prognostic or diagnostic markers or therapeutic targets, we selected SAA2, CFB and IL34 for their presence in the signatures and because they also represent potential soluble markers that can be analyzed in the blood. We therefore carried-out a more detailed analysis.

#### Serum Amyloid A2 (SAA2)

SAA2 is an acute phase protein related to SAA1, which was previously linked to metastasis *(Hansen et al, 2015)* increased in lung cell lines (fig. S8A) and present in the Lung signature. *In silico* analysis of the TCGA KIRC database indicated that *SAA2* was a very strong predictor of OS and DFS (fig. S8, B and C). Fhe analysis was also done for the M0 and M1 subgroups (fig. S8, D to G). Analysis of the UroCCR patient cohort confirmed the effect on OS and DFS (fig. S8, H and I). Tumors from patients with the highest Fuhrman Grade, had a significantly increased SAA2 expression compared to all other grades (fig. S8J). We used grade-matched plasma samples from patients with and without metastases, collected before primary tumor surgery (fig. S8K). Patients with metastases had higher plasma levels of SAA2. When patients were divided into two groups of equivalent size, the group with higher SAA2 levels had a significantly shorter DFS (fig. S8L). A second set of plasma samples, collected in the weeks following surgery for removal of the primary tumor, was tested for SAA2 (fig. S8M). In this case, patients with higher expression had shorter OS. Hence, circulating SAA2 levels appear as an indicator of metastatic progression that deserves to be evaluated at diagnosis.

We next used plasma samples from metastatic patients before receiving a first cycle of sunitinib or bevacizumab (SUVEGIL and TORAVA clinical trials). Patients treated with sunitinib only and stratified according to low and high SAA2 levels, had a spectacular better OS and progression-free survival (PFS) when belonging to the SAA2 low group (cut-off of 269 µg/ml) (Fig. 5, A and B). When patients treated with sunitinib and bevacizumab were analyzed together, the PFS was of limited significance (borderline p-value of 0.0507) (Fig. 5C). The median of PFS for SAA2high patients was of 5.35 month versus 16.17 month for the SAA2low group. Thus, determining SAA2 plasma levels could be a useful measure for deciding a treatment strategy in RCC.

**Fig. 5.**
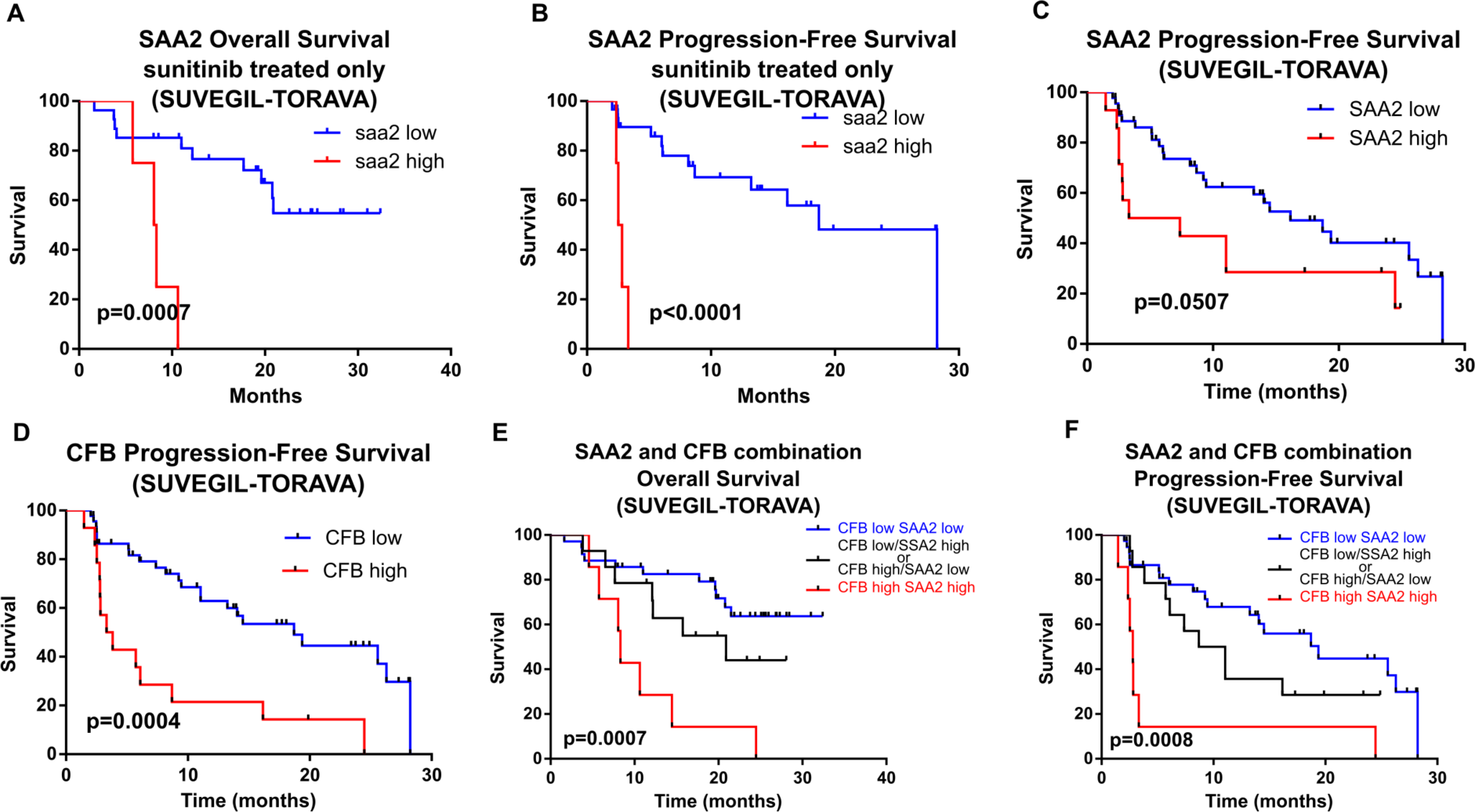
Clinical relevance of SAA2 and CFB after anti-angiogenic treatment (SUVEGIL-TORAVA cohorts). (**A** and **B**) Correlation between plasmatic SAA2 levels at diagnosis and survival (OS and PFS) in patients after sunitinib treatment (plasmatic level at the diagnosis less or greater than a cut-off for SAA2 (269 µg/ml)[OS: HR(log-rank)=5.557; PFS: HR(log-rank)=7.669. (**C**) Correlation between plasmatic SAA2 levels at diagnosis and PFS in patients after sunitinib or bevacizumab treatment (plasmatic level at the diagnosis less or greater than a third quartile cut-off for SAA2 (269 µg/ml; HR(log-rank)=1.987). (**D**) Correlation between plasmatic CFB levels at diagnosis and PFS in patients after sunitinib or bevacizumab treatment (plasmatic level at the diagnosis less or greater than a third quartile cut-off for CFB (310µg/ml; HR(log-rank)=3.113) (**E** and **F**) PFS (**E**) and OS (**F**) patients treated with either Sunitinib of bevacizumab and stratified according to plasma levels of both SAA2 and CFB. Three subgroups were identified *i)* CFB low and SAA2 low, *ii)* CFB low and SAA2 high or CFB high and SAA2 low, *iii)* CFB high and SAA2 high (Low-low vs high-high: OS HR(log-rank)=5.086; PFS HR(log-rank)=4.196).

#### Complement factor-B (CFB)

CFB was most strongly upregulated in the “Lung” and to a lesser extent in the “Tail” group, both considered to recapitulate features of metastasis (fig. S9, A and B) and was also present in the Lung signature. TCGA analysis in ccRCC showed that *CFB* expression is correlated in primary tumors with shortened DFS and OS (fig. S9, C and D). We also performed the analysis in the M0 and M1 subgroups (fig. S9, E to H). Using samples and data from the UroCCR cohort, we demonstrated that *CFB* was overexpressed in the tumor tissue versus the adjacent kidney at the mRNA level (fig. S9I), and that increased expression correlated with reduced DFS and OS, consistent with the results obtained with the TCGA cohort (fig. S9, J and K). As for SAA2, CFB can be measured in the blood. For this purpose, we used UroCCR plasma samples collected from patients either before surgery (primary tumor intact) or in the following weeks after surgery (no primary tumor present but metastases *in situ* possible). Before surgery, a trend was observed without reaching significance whereas after surgery patients with metastases had higher plasma CFB levels compared to patients without metastases (fig. S9, L and M). This suggests that circulating CFB measurement may be useful as a blood-born marker of metastasis in the follow-up after surgical tumor removal. As for SAA2, CFB plasma levels were tested in patients with metastases before the first cycle treatment with sunitinib or bevacizumab (SUVEGIL and TORAVA clinical trials). Patients whose levels were high (cut-off 310 µg/ml) had faster disease progression compared to patients whose levels were low (high CFB, 3.58 month; low CFB, 18.7month, p=0.0004) (Fig. 5D).

We then grouped the significance of testing SAA2 and CFB plasmatic levels (Fig. 5, E and F). Three different groups with different survival can be identified: group 1 (CFB low+SAA2 low, PFS: 19.37 months, OS: NR), group 2 (CFB high SAA2 low or CFB low SAA2 high, PFS: 9.87 months, OS: 20.9 months), group 3 (CFB high Saa2 high, PFS: 2.8 month, OS: 8.33 months). Group 1 had the best survival rate while group 3 had the worst. Group 2 had intermediate survival outcome. Thus, the combined analysis of these two markers is a powerful predictor of patient outcome following anti-angiogenic treatment with sunitinib or bevacizumab.

### Expression, clinical relevance and functional analysis of Interleukin-34

*IL34* expression was upregulated in all cell line groups (Fig. 6A) and is present in the “Kidney” and the KTL signature. We determined in an independent experiment that expression was increased in both in primary tumor cells and lung metastatic cells (Fig. 6,B). Analysis by ELISA demonstrated that higher amounts of IL34 protein were secreted into the culture medium by Passage 6 in “Kidney”, “Tail” and “Lungs” cells when compared to parental cells (Fig. 6C). *In silico* analysis in TCGA demonstrated that higher IL34 mRNA expression correlated with shortened OS and DFS (Fig. 6, D and E). This was also analyzed in M0 and M1 patients where IL34 expression was significantly correlated to OS and DFS in M0 patients (fig. S10, A to D). Furthermore, *IL34* expression was also correlated to tumor grade in TCGA (Fig. 6F). In the UroCCR cohort, IL34 mRNA expression was higher in tumors compared to healthy tissue (Fig. 6G). A Tissue MicroArray (TMA) from the same set of samples was stained for IL34 and graded and scored by a pathologist (Fig. 6H). Immunohistochemical staining correlated with Fuhrman grade, and was predictive of DFS (Fig. 6, I and J). We also tested a limited number of plasma samples from a different cohort of patients undergoing treatment for metastases with the receptor tyrosine kinase inhibitor sunitinib (SUVEGIL an TORAVA clinical trial) (fig. S10E). In these conditions, a subset of patients (6/20) showed a marked spike in plasma IL34 following the first cycle of treatment. The sample size is too small to conclude a link between this spike and clinical parameters such as treatment response, and requires further investigation. However, in a xenograft mouse model (human 786-0 ccRCC cell line implanted s.c. in immunodeficient mice), treated with sunitinib, *IL34* mRNA expression was increased in tumor cells (fig. S10F, left panel).

**Fig. 6.**
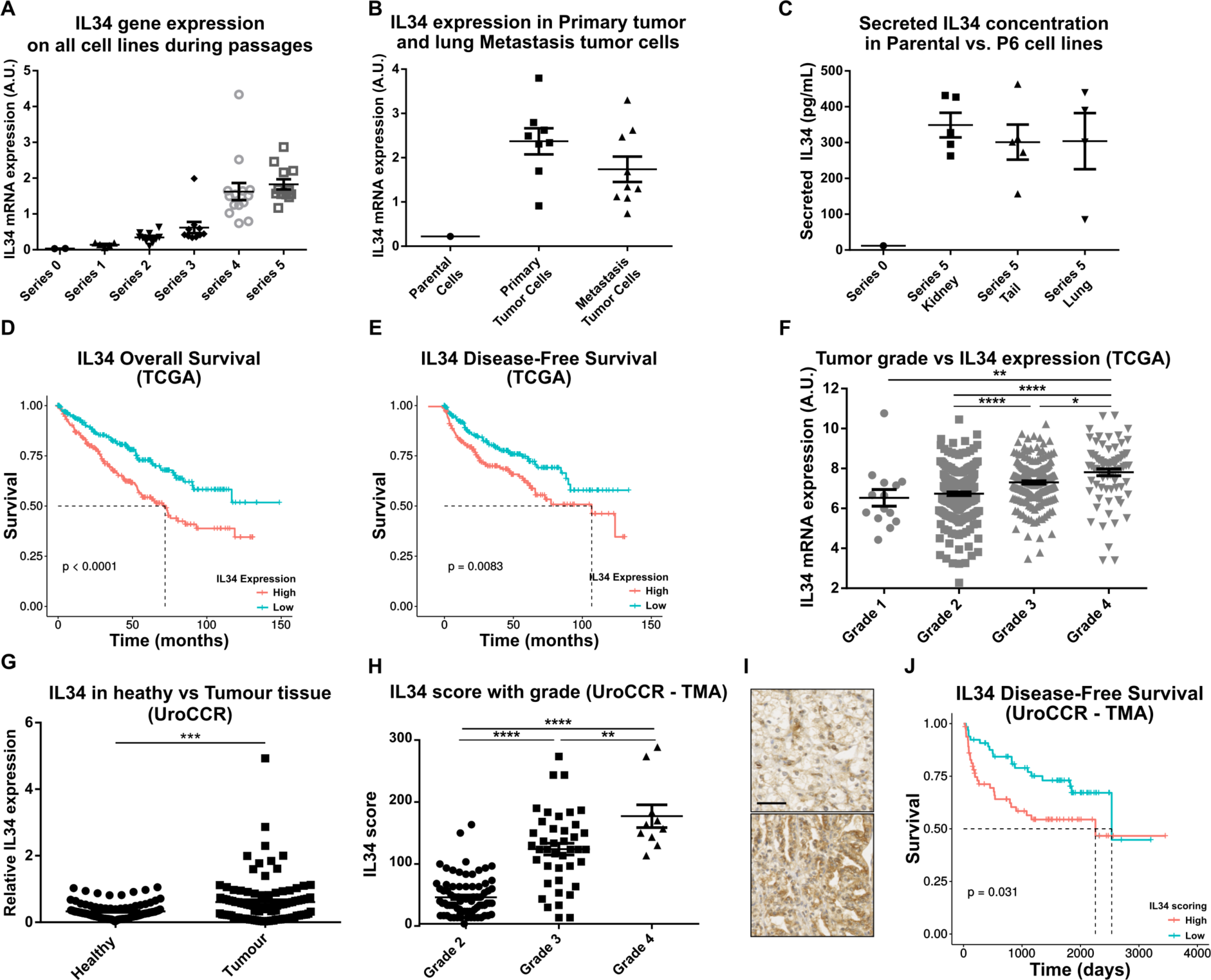
Analysis of IL34 expression and its significance for RCC progression. (**A**) Expression analyses of IL34 gene expression in amplified RENCA cell lines (Kidney, Tail and Lung) where IL34 was strongly upregulated compared to Parental cells (Series 0). Of note, the increase of expression is cell-passage dependent and, thus, related to progression toward a more aggressive phenotype. Graph shows mean ± SEM. (**B**) IL34 expression in primary tumor cells and in tumor cells from lung metastasis compared to the parental cell line. (**C**) ELISA assay of IL34 in supernatants in late passaged (S5) cell lines compared to control (Parental S0) of 1 million cells. Graph shows mean ± SEM. (**D**) Correlation of IL34 expression to overall survival (OS) in patients from TCGA KIRC cohort ((HR(log-rank)=1.857 (1.38-2.498)). (**E**) IL 34 expression correlated to Disease Free Survival (DFS) in patients from TCGA KIRC cohort. ((HR(log-rank)=2.056 (1.315-3.214)) (**F**) Correlation of Fuhrman grade with IL34 gene expression in the TCGA KIRC cohort. (**G**) qPCR analyses of IL34 expression in healthy kidney and in tumors from patients (UroCCR). Results as mean and SEM. Analysis by two-tailed paired t-test. P=0.0006. (**H**) Evaluations of IL34 expression in TMA from RCC patients (UroCCR cohort) (Scoring by a pathologist, D. Ambrosetti). ANOVA followed with tukey’s multiple comparisons. **p<0.01; ****p<0.0001. (**I**) Histological staining of IL34 in tumors from patients (UroCCR). Scale bar = 50µm. (**J**) Disease Free Survival (DFS) from the TMA IL34 scoring (UroCCR Cohort)HR(log-rank)=1.878 (1.056-3.342).

Stromal mouse *IL34* showed a similar trend towards increased expression indicative of a similar, though weaker, host response (fig. S10F, right panel). To further study the role of IL34 in RCC, we overexpressed IL34 (variant 1) in RENCA-GFP cells, using a lentiviral vector and analyzed *in vivo* tumor development (fig. S11, A to C). Tumors generated with IL34-overexpression versus control cells did not show significant altered size, and lifespan of mice was not changed (fig. S11D). IL34 expression was validated in tumors at the mRNA and protein level (fig. S11, E and F). We next hypothesized that IL34 overexpression could alter the balance of Type 1 and 2 macrophages within the tumor and analyzed the levels of various macrophage markers in IL34-overexpressing versus control primary tumor lysates. IL34-overexpressing tumors contained higher levels of the total macrophage markers *F4/80* and *CSF1R* compared to control tumors indicating an increased macrophage number (Fig. 7A). No significant difference was detected in levels of the Type 1 markers *IL1β* and *IL12a* (Fig. 7B), whereas the Type 2 markers *MMR* and *ARG1* were increased in IL34-overexpressing tumors (Fig. 7C). Furthermore, *MMR* and CSF1R mRNA levels correlated positively with the level of *IL34* in primary tumors (Fig. 7D and fig. S11G). Increased MMR expression in IL34 overexpressing tumors was validated at a protein level by Western Blot (Fig. 7E). Importantly, the level of CSF1, a twin cytokine to IL34 and also involved in macrophage attraction and differentiation, was not altered (fig. S11H). Furthermore, an analysis of MMR-stained tumor sections revealed an increased density of MMR-positive cells in the primary tumor (Fig. 7F).

**Fig. 7.**
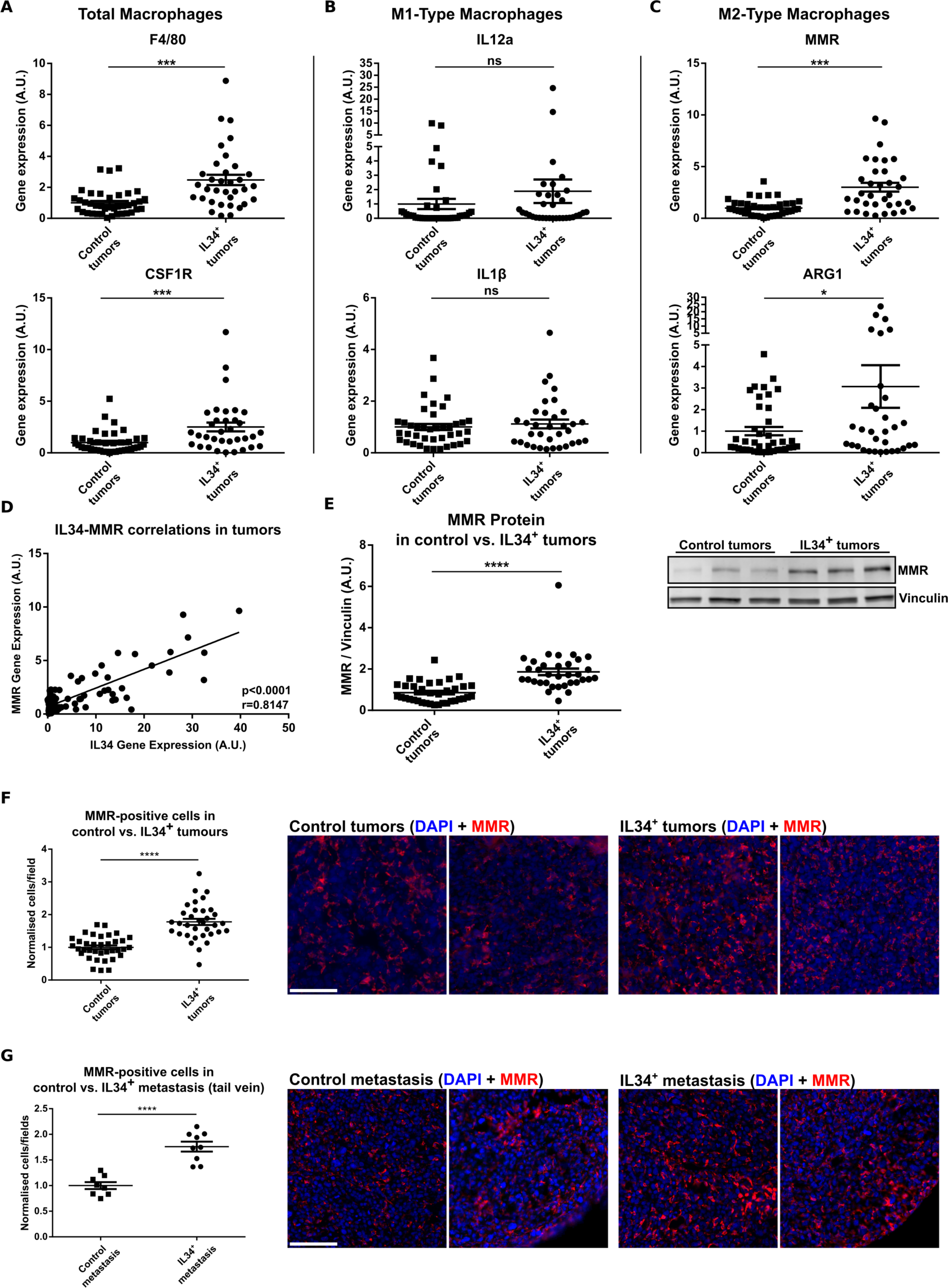
Analysis of IL34 overexpression on macrophages. Expression analysis conducted in orthotopic implanted tumors generated from IL34-overexpressing RENCA cells. Expression of markers in IL34-overexpressing tumors for (**A**) total macrophages (F4/80 and CSF1R), (**B**) M1-type macrophages (IL12a and IL1β) and (**C**) M2-type macrophages (MMR and ARG1) compared to control tumors. Graph show mean ± SEM. T-test. *p<0.05; ***p<0.001. (**D**) Correlation relationship between IL34 and MMR (type 2 macrophage marker). Statistical test: Spearman correlation test. (**E**) Semi-quantitative analysis of MMR protein expression in mouse tumors generated from IL34-overexpressing and control RENCA cells from western blots (Quantification of band intensities compared to vinculin and using a reference sample). On the right, a representative WB of MMR in IL34-overexpressing and control tumors. T-test. ****p<0.0001. (**F**) Quantification of MMR-positive cells in the kidney tumors. On the right, representative images of primary tumors stained by IF using an anti-MMR fluorescent antibody (red) and DAPI (blue) for DNA counterstaining. Scale bar, 100 µm. T-test. ****p<0.0001. (**G**) Quantification of MMR-positive cells in lung metastasis after tail vein injection. On the right, representative images of lung metastases stained by IF using an anti-MMR fluorescent antibody (red) and DAPI (blue) for DNA counterstaining. Scale bar, 100 µm. T-test. ****p<0.0001.

Analyses of expression in metastases are limited by the relatively low number of metastases using the orthotopic model. We, therefore, used intravenous tail vein injection of IL34 overexpressing tumor cells. Analysis of these tumors did not show a significant increase in the metastasis area compared to control (fig. S11I), but an increased density of MMR-positive macrophages (Fig. 7G), similarly to orthotopic primary tumor results, was detected.

Taken together, these data suggest that IL34 overexpression mediates an influx of macrophages with a polarization towards the type 2 phenotype. Since IL34 overexpression alone seems to be insufficient to increase tumor size or speed of progression, we deleted IL34 in RENCA cells with two different Crispr/cas9 constructs (CrisprIL34_1a, CrisprIL34-1c) and used a targeting LacZ vector as a control (Crispr_LacZ). Interleukin-34 deletion was validated by ELISA (fig. S12A). When implanted orthotopically, growth of IL-34-deleted primary tumors was strongly inhibited (Fig. 8A), albeit *in vitro* cell proliferation was not affected (fig. S12B). We, next, used magnetic resonance imaging (MRI) to track primary tumor volume dynamics, and appearance of metastases in the orthotopic model for one of the IL34-deleted cell lines (≥5 mice/group). Survival was significantly increased for the IL34-deleted cell line (Fig. 8B) and metastasis appearance was delayed (Fig. 8C). Tumor development and spread was then mathematically modeled using ordinary differential equations (ODE, see: supplementary Methods) (Fig. 8D). In the IL34 Crispr group, 3 mice did not develop primary tumors and were therefore not included in the ODE modeling. Furthermore, two of these mice did not develop metastasis and were also excluded. Consequently, the prediction of primary tumor volume and of metastasis number for the IL34 Crispr group shown in Fig. 8D is overestimated, but nevertheless demonstrated slower primary tumor growth and metastatic spread. The data were fitted using a population approach by means of a non-linear mixed effect model (see: supplementary Methods). This approach allows individual modeling of growth and dissemination kinetics, while also quantifying inter-animal variability and group effects. It is particularly adapted to sparse data and strengthens the statistical power. The data fitted reasonably well the mathematical model, both at the population (Fig. 8D) and individual level (fig. S12C) which is composed of two parameters: a growth coefficient α and a dissemination coefficient µ. A negative group effect for IL34 KO was observed for both α (p=0.0615) and µ (p=0.053) (table S3).

**Fig. 8.**
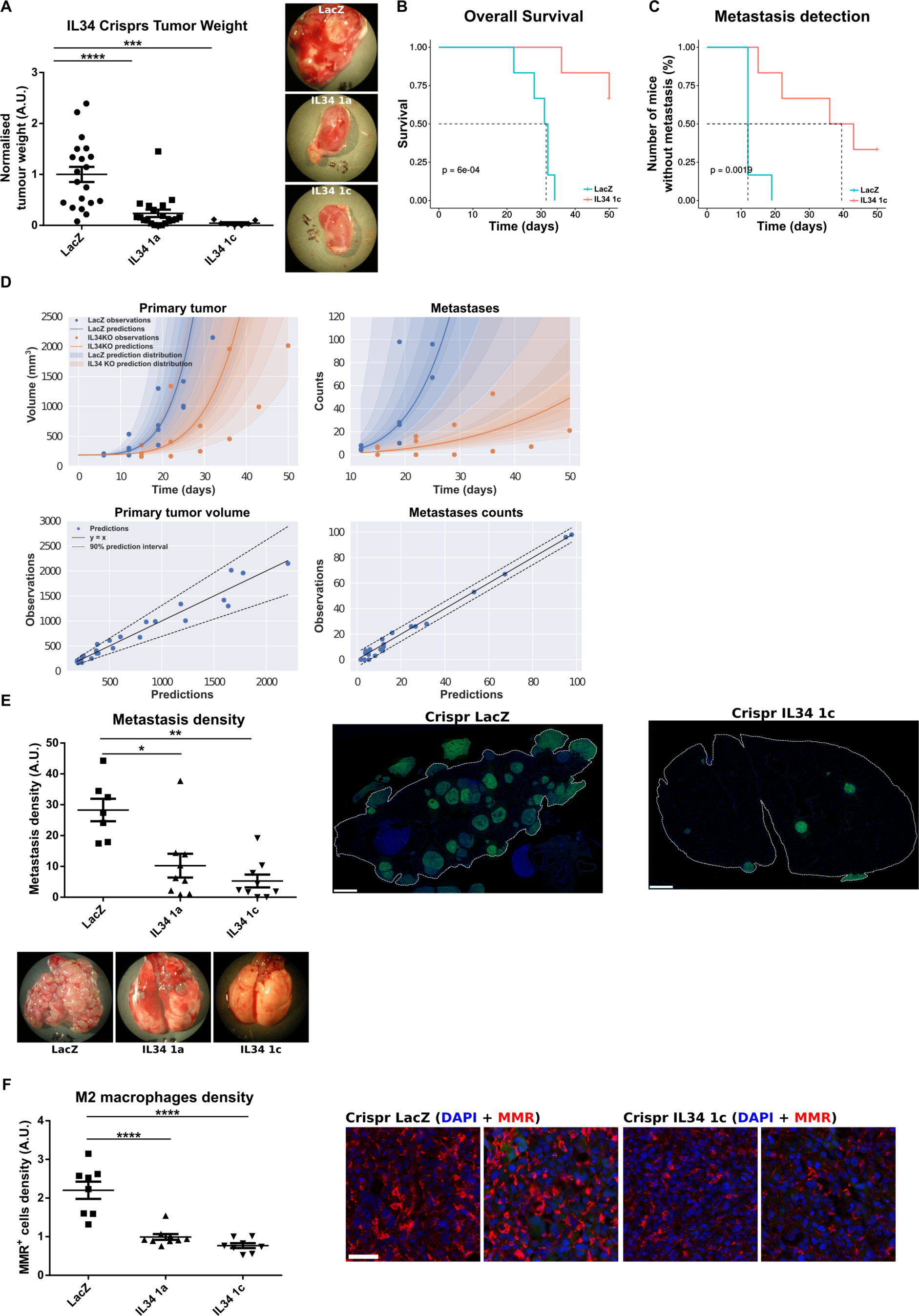
**Effect of IL34 silencing *in*** *vivo*. (**A**) Weight from kidney bearing tumors after orthotopic implanted IL34 knock-out (Crispr-Cas9) or control (LacZ) cells represented as mean+-SEM. The pictures on the right shows images of implanted tumors. ANOVA followed by Dunnett’s multiple comparisons test. ***p>0.001; ****p<0.0001. (**B**) Mouse survival (HR(log-rank)=8.09 (1.7-38.497)) and (**C**) number of mice without lung metastasis (HR(log-rank)=3.839 (0.967-15.244)). (**D**) Mathematical modeling issued of tumors from sub-capsular implant of IL 34 Crispr or control (LacZ) cells. Volume of primary tumors and the number of lung metastasis were quantified by MRI and fitted to generate simulations for tumor growth and metastasis formation, using a population mixed-effects approach. (**E**) Metastasis density after tail-vein injections of IL34-silenced and control (LacZ) cells. The graph shows the means of the metastasis area in lung sections relative to the tissue area ± SEM. Examples of macroscopic images of lungs, and lung metastasis from GFP+ RENCA cells are shown. Statistical analysis: ANOVA followed by Dunnett’s multiple comparisons test. *p<0.05; **p<0.01. (**F**) Quantification of MMR-positive cells in lung metastasis after the tail vein injection. Representative images of IF staining for MMR (red) and DAPI(blue) are shown. Scale bar = 100µm. Statistics: ANOVA followed by Dunnett’s multiple comparisons test. ****p<0.0001

To examine macrophage density in the lungs after IL34 deletion, we used the tail vein metastasis model since this model is more efficient in generating lung metastasis. Metastasis formation was drastically reduced using the two CrisprIL34 cell lines (Fig. 8E). We used lung sections stained for GFP to calculate tumor area/Lung area, and MMR staining to calculate macrophage density. Within the metastases, MMR+ cell density was reduced in IL34-deleted tumors (Fig. 8F).

## DISCUSSION

The usual strategy in translational research is to start from human samples, to identify molecular markers and gene networks and then to functionally validate them *in vitro* and in animal models. We devised herein a completely opposite strategy (from “mouse to man”) starting from the animal model and used functional genomics, imaging, clinical data and computational approaches. We took advantage of a syngeneic mouse model and developed it further to unravel mechanisms of RCC development and metastasis formation, and to identify novel therapeutic targets and biomarkers.

The strengths of our model are the following. The model is based on iterative cycles of injection using 3 different injection modes capturing local tumor development (“Kidney” group), metastasis formation from the primary tumor involving all steps of metastasis formation (“Lung” group), or only the last steps of metastasis formation (“Tail” group). With increasing cycles, a shortening of the mice’s lifespan was observed. The model is, furthermore, a syngeneic mouse model which leaves the adaptive immune system intact. Another important strength of our model is that the 3 different groups yield very distinct transcriptomic and methylome clustering and signatures, which are clinically relevant.

The model recapitulate critical steps of RCC formation and spread and is robust and reproducible *(Tracz et al, 2014)*. Our aim was to uncover the molecular changes that are occurring with increasing aggressiveness and tumor specialization and not the genetic characteristics of the tumor itself, for which this RENCA model is well suited. This is different from human ccRCC models including implantation of cell lines or patient derived xenografts (PDX) which require longer time to develop in an immunocompromised context and are mainly focused on primary tumor growth. In addition, no suitable genetically engineered mouse model (GEMM) for RCC that recapitulates with fidelity the metastatic process exists to date *(Di Martino et al, 2018; Hillman, 2002; Schueler et al, 2018; Patel et al, 2019)*. It is true that around 10 different GEMMs for RCC have been developed *(Hou & Ji, 2018)* such as the MYC activation, Vhl del Cdkn2adel, Ink4a/Arf del model *(Bailey et al, 2017)*, the Bap1 and Pbrm1 inactivation model *(Gu et al, 2017)*. Only the Vhl del Ink4a/Arf del model produces some metastasis but only in the liver, which is not suitable for our study. Thus, even if our model does not strictly reflect the exact characteristics of a human RCC tumor, it, nevertheless, recapitulates critical steps of tumor development and spread of a renal carcinoma.

In the sequentially amplified cell lines, RNA expression and methylation analysis demonstrated distinct clustering for the 3 different injection modes. DNA sequencing did not show clonal variations based on chromosomal variability, which indicate that the phenotypic changes were epigenetically regulated. Transcriptomic analysis led to the identification of specific gene signatures for each injection cycle which were predictors of overall, disease-free or progression-free survival in RCC as based on TCGA analysis and on our RCC cohorts. Importantly, some of the signatures, especially the Lung signature, are stronger predictors than current predictors in clinical use such as Fuhrman Grade or clinical stage. Beuselink et al determined four ccRCC signatures (ccrcc1-4 signatures) from the transcriptome of metastatic patient treated sunitinib *(Beuselinck et al, 2015)*. The HR reported in their article is lower than in our study. In a follow-up study by this team, Hazard ratios (HRs) were calculated in a patient cohort treated with pazopanib which is different to our study *(Verbiest et al, 2018)*. Recently, Ricketts et al *(Ricketts et al, 2018)* reported a comprehensive molecular characterization using the TCGA database where they compared the 3 types of RCC (ccRCC, papillary RCC and chromophobe RCC). For ccRCC, the signatures were related to increased ribose metabolism pathway and to Th2 immune profile. However our study is very different, because starting from an animal model we specifically focused our comparative translational analysis on the ccRCC subtype and, thus, the results cannot be compared, albeit their study also revealed immunology-related gene expression. Furthermore, a recent study reported tracking of ccRCC evolution at the genomic level and demonstrated that metastatic competence was afforded by chromosomal complexity with loss of 9p as a selective event for metastasis and patient survival *(Turajlic et al, 2018)*. Our study did not include an analysis of the chromosomal events, since our animal model did not reveal chromosomal alterations and, thus, we specifically focused on modifications of the transcriptome and the methylome.

Methylome analysis from the S5 series revealed that only 29 % of 33095 CpGs differentially methylated genes were associated with a known gene. Only a restricted number of genes found in this analysis was contained in the transcriptome signatures which include *IL34*, *Krt17*, *RIK/MAGEC2/3*, and *SAMSN1*. Thus, despite the excellent clustering only a restricted number of genes are in common. When GO terms are crossed between transcriptomics and methylome data sets, they also, only, converge partially to a common set of processes. The reason for this resides in the fact that many methylation sites are found within intronic sequences, the role of which is not really understood, but which may have major regulatory functions. For IL34, three CpGs were identified as part of enhancer regions with two of those found in the gene body and predictive for OS which points to a possible epigenetic regulation of IL34 (see for more on IL34 later in discussion).

We further selected, for a more detailed investigation, SAA2, CFB and IL34 which we showed to be predictive of patient outcome in the TCGA database and verified in a second local patient cohort.

SAA2, an acute phase protein related to SAA1, was found to be a strong predictor of OS and DFS in the TCGA KIRC database globally and also for the M0 and M1 subgroups. In patients with the highest Fuhrman Grade, significantly increased SAA2 expression was observed. Furthermore, analysis of plasma samples from patients with metastases before or after surgery showed higher plasma levels of SAA2 and worse OS. Analysis of samples from the clinical trials evidences SAA2 as an excellent predictive biomarker especially in the sunitinib treated patient cohort. Serum amyloid A has been previously reported to be prognostic marker in RCC *(Wood et al, 2010; Vermaat et al, 2009, 2012)*. In the Vermaat et al study *(Vermaat et al, 2012, 2009)*, there is no distinction between the four SAA variants (SAA1-4) contrary to our study where we clearly identified SAA2. In addition, in their first article *(Vermaat et al, 2009)*, most of the patients included in the study were treated with interferon as first line treatment and not with anti-angiogenic drugs currently in clinical use, and in this case PFS was not analyzed and only the combination with apolipoprotein A2 (ApoA2) was predictive for OS. Their second study *(Vermaat et al, 2012)*, which demonstrated highly significant predictive value for SAA in RCC metastatic patients treated by tyrosine-kinase inhibitors (TKI), is in agreement with our data, albeit they did not specify the SAA variant.

Another molecule of interest is complement factor B (CFB) which was upregulated in “Lung” and “Tail” cell lines. Complement factor B expression correlated with survival and metastasis in the TCGA data set and in the UroCCR cohort. Plasma measurements showed that, similarly to SAA2, patients with metastases had higher CFB plasma levels compared to patients without metastases. This remained the case whether the samples were taken before or after surgery, suggesting that CFB may be a useful blood borne marker of metastasis. Like SAA2, CFB plasma levels were tested in patients with metastasis treated with antiangiogenic drugs, before and after a first cycle treatment with sunitinib. Patients whose CFB levels were increased following treatment had faster disease progression and shortened survival compared to patients whose level were decreased. Interestingly, the combination of SAA2 and CFB plasma level measurements produced a more powerful analysis than either marker alone. This suggests that a combined analysis of these two markers represent a powerful tool for predicting patient outcome following anti-angiogenic treatment. Interestingly, CFB is a prognostic preoperative marker in pancreatic carcinoma which outperformed CA19-9 and CEA *(Kim et al, 2019)*. However, the combination of CFB and SAA2 was not tested in this study.

One of the most interesting genes in our study was Interleukin-34 (IL34), a secreted cytokine, which binds to CSF1R *(Lin et al, 2008)*, receptor-type protein-tyrosine phosphatase ζ (PTP-ζ) *(Nandi et al, 2013)* and Syndecan-1 *(Segaliny et al, 2015)*. It is expressed in several organs, including in tubular cells of the kidney *(Baek et al, 2015)*. IL34 supports the growth and survival of primary human monocytes and promotes their differentiation into mature immunosuppressive macrophages *(Foucher et al, 2013)*. IL34 has been linked with autoimmune and inflammatory diseases such as rheumatoid arthritis *(Foucher et al, 2013; Boulakirba et al, 2018)*, hepatitis C infection *(Chang et al, 2015)*, lupus nephritis *(Preisser et al, 2014)*, inflammatory bowel disease *(Wada et al, 2019)*. It has been linked to chronic kidney injury and kidney graft rejection *(Baek et al, 2015)*, and to cancers including osteosarcoma *(Ségaliny et al, 2015)* or lung cancer *(Baghdadi et al, 2018)*.

In our study, interleukin-34 was upregulated in the aggressive mouse cell lines, suggesting that it could contribute to different stages of RCC progression, acting as a key driver/regulator of aggressiveness. We showed that its expression in RCC correlated with increased Fuhrman grade, and shortened survival and faster metastatic progression, using two cohorts of patients, and RNA and immunohistochemistry data. We also provided evidence that IL34 is detected in the plasma of a subset of patients treated with sunitinib, the first line treatment for metastatic RCC.

For functional analysis, we focused on IL34 since IL34 was a common driver to all experimental groups and since our data also indicated a potential epigenetic regulation of this gene with two potential methylation sites predictive for OS and PFS. We first overexpressed IL34 in RCC and determined the effect on macrophages and tumor development in the mouse model. Our studies showed that mouse tumors overexpressing IL34 had an altered pool of tumor-associated macrophages with an increase in type 2 pro-tumorigenic macrophages while type 1 macrophages were not significantly altered. It is known that type 2 macrophages are associated with worsened progression in multiple cancer types *(Baghdadi et al, 2018)*.

Although IL34 overexpression alone was not sufficient to further increase tumor progression in our model, CrisprCas9-mediated deletion of IL34 strongly inhibited primary tumor formation and metastasis. However, *in vitro* proliferation in IL34 KO cells was not modified, suggesting that this effect is microenvironment-dependent. Mouse survival time was increased, and MRI analysis showed reduced numbers and slowed development of metastases which was also mathematically modeled. Importantly, RENCA cells do express CSF1, but our experiments showed that the loss of IL34 cannot be compensated by the presence of CSF1. Our data also point to the fact that IL34 is already required at an early stage of tumor development. Taken together, these studies demonstrate a previously unknown role for IL34 in progression of RCC and suggest it as an important potential target. IL34 has been also shown to promote tumor progression and metastasis in other tumor types such as osteosarcoma *(Guadagno et al, 2018; Franklin et al, 2014; Sica et al, 2014; Comito et al, 2014; Funada et al; Kurahara et al, 2011; Ryder et al, 2008)*. The effect of IL34 and CSF1 blockade was recently explored using specific blocking antibodies *(Lin et al, 2019)*. Mouse colon carcinoma (MC38 cell line) development was only weakly inhibited by both IL34 and CSF1 blockade but only CSF1 blockade had an effect on TAM homeostasis and CD4^+^ T cell accumulation which is very different from our data.

All in all, our work demonstrates that a multilayered systems approach, from mouse to man as described in this work, generates many meaningful results that are not only important for the understanding of tumor biology but which are also highly relevant for clinical translation.

## Methods

### Mouse tumor and metastasis models

The Renca murine renal cancer cells, and the human 786-0 cancer cells were maintained in Roswell Park Memorial Institute (RPMI) 1640 medium supplemented with 10% feral bovine serum (FBS) and 1% penicillin/streptomycin (Complete Medium), and were incubated at 37°C, 5% CO_2_ in an incubator.

For sub-capsular implantations of Renca cells, 1*10^5^ cells were injected under the left kidney capsule of 6-week-old female BALB/c mice (Charles River Laboratories).

For intravenous injections of Renca cells, 5*10^6^ cells were injected into the caudal vein (tail vein) of 6-week-old female BALB/c mice. When the endpoints defined by the approved protocols were reached, mice were sacrificed and tumor tissues and lungs were collected. When a first mouse from a group was determined as having reached an endpoint, all mice from that group were sacrificed. Mice were housed in the animal facility of Bordeaux University (Animalerie Mutualisée, Université de Bordeaux, France).

Sub-cutaneous implantations of 786-O cells, tissue dissociation, tumor cell purification and tissue processing are described in Supplementary Materials and Methods.

All animal experiments were approved by the “Ministère de l’Enseignement Supérieur, de la Recherche et de l’Innovation (MESRI)” (authorizations numbers 2016072015478042; 2015110618597936 and 2015070315335217), and were carried out in accordance with the approved protocols.

### Low-coverage whole-genome sequencing

Genomic DNA was isolated use the Dneasy Blood and Tissue Kit (Qiagen, 69504), according to the manufacturer’s protocols. 13 samples were subjected to low-coverage whole-genome sequencing as described in supplementary Materials (Method section).

### Transcriptomic data generation and analysis

Total RNA was extracted using the RNeasy Plus Mini Kit (Qiagen, #74134), according to the manufacturer’s protocols. Analyses of transcription sets, signature computation and enrichment analysis were done using R studio (R v3.5.2 *(Carter et al, 2012)*, R studio v1.1.463 *(R Foundation for Statistical Computing., 2018)*).

### Biomarker discovery

***(i) Differential expression between parental cell line (series 0, S0) and series 5 (S5):*** To select genes that present the strongest difference in expression between parental cell line and series 5 experiments, we used a z-score approach. First, we compute the log fold (logFC) change for each gene. Second, we compute the mean and standard deviation of the logFC and a z-score for all genes. A gene was considered as differentially expressed if the absolute value of its z-score was ≥ 2.58 and if its logFC is ≤ −1 (down expressed) or ≥ 2 (up expressed). ***(ii) Progressive expression pattern through series:*** To capture genes that present a progressive expression through series, we compute mean values of expression for each series. A gene will be considered progressive if its mean values between replicates for a given series are strictly increasing (up) or decreasing (down) through series 0 to 5. As the S1 of Lung group had only one measure, it was not usedfor the (determination of the (progressive quality of genes. ***(iii) Differentially expressed and progressive genes.*** A gene was considered differentially progressively expressed through series if it was in the intersection of the gene sets selected in steps (i) and (ii) with the additional consistency constraint that the expression should be in the same direction (up or down).

### Biomarker validation

Only genes whose overexpression was predictive of a bad prognostic were conserved. (*i) Selection of clinically relevant genes:* In order to conserve genes that are relevant in human kidney cancer, we used the TCGA KIRC cohort. For each gene, we fitted a Cox proportional hazard regression model based on overall survival (OS) time and disease-free survival (DFS). A gene was conserved if its cox proportional hazard ratio (HR) was in accordance with the differential expression (HR ≥ 1 for up regulated genes and HR ≤ 1 for down regulated genes) and if the FDR adjusted p-value of its log-rank test was ≤ 0.01 for OS and/or DFS. (*ii) Relevance of the association of selected genes:* To measure the clinical relevance of the resulting signature and check their prognostic performance against random signatures, we used SigCheck R package (v 2.14.0). To separate samples into groups, we computed a score corresponding to the mean value of all the expression values of genes for each sample ((scoreMethod=“High” in the sigCheck function). Patients were then classified in accordance to their score and splited in 3 groups using a log-rank test followed by computation of the associated log-rank HR (survival v2.44-1.1 and survminer v0.4.3 R packages). We then compared the performance of our previously defined signatures with 1000 signatures composed of the same number of randomly selected genes. TCGA Kidney Renal Clear Cell Carcinoma (KIRC) HiSeqV2 data were downloaded from XenaBrowser *(Rstudio Team, 2016)*. We chose log2(x+1) transformed RSEM normalized count (version 2017-10-13) as recommended on its web site (https://xenabrowser.net/; page: «dataset:gene expression RNAseq-IlluminaHiSeq percentile»). We removed genes where there were more than 90% of samples that had null values resulting in 19186 genes. Complementary associated clinical data were download from cbioportal (https://www.cbioportal.org/). Conversion of *Mus Musculus* to Human gene name was done by downloading conversion table from the Biomart website (https://www.ensembl.org *(Goldman et al, 2019)*). Methylomics data generation and analysis is described in supplementary Material (Method section).

### Enrichment analyses

To perform enrichment analyses we used hypergeometric test (enricher function of Cluster Profiler R package *(Fishilevich et al, 2017)*; v3.10.1) with go_terms.mgi download on Mouse Genome Database (MGD) at the Mouse Genome Informatics website (URL: http://www.informatics.jax.org *(Yu et al, 2012)* (04, 2019). Enrichment analysis was done for following gene sets: (i) transcriptomics data: gene set related to PC1, gene set related to PC2 and gene sets related to PC1 and PC2 (universe was setting to the set of genes which are measured in transcriptomic experiments) (ii) methylomics data: gene sets related to PC1 and PC2.

### Immunochemistry and immunofluorescence

This was done as previously described *(Daubon et al, 2019)* and is detailed in supplementary Methods.

### Generation of GFP, IL34, overexpressing cells and crispr/cas9 induced IL34 knock-down cells

This is indicated in Supplementary Methods.

### qPCR gene expression analysis

Total RNA was extracted using the RNeasy Plus Mini Kit (Qiagen, #74134), according to the manufacturer’s protocols. 1µg of total RNA was reverse-transcribed into complementary DNA (cDNA) using the high-capacity cDNA reverse transcription kit (Applied Biosystems, 4368814). The resulting cDNA were amplified using specific primers for the genes of interest. For RT-qPCR, Taqman (Eurobio Scientific, EurobioProbe) or SybrGreen (Eurobio Scientific, EurobioGreen) technologies were used. HPRT was used as internal control. Primer sequences are indicated in Supplementary Materials (Method section).

### Western Blot analysis

Western blots were done as described *(Daubon et al, 2019)* and as indicated in Supplementary Material (Method section). A fluorescent secondary fluorescent antibody (see table) was used as detection system. Membranes were scanned with an infrared imaging system (Odyssey, Li-Cor Biosciences, Nebraska, US). Densitometry analysis were performed using Image Studio Lite 4,0 Software (Li-Cor).

### ELISA

ELISA were performed according to the manufacturer’s protocols: human IL34 (R&D systems, D3400), human SAA2 (DLDEVELOP, DL-SAA2-Hu-96T), Human CFB (abcam, ab137973).

### Human patient samples

Patient samples (tumor tissue and plasma) from the UroCCR cohort were used with associated clinical data (clinicaltrial.gov, NCT03293563). Eligible patients for SUVEGIL and TORAVA trials were at least 18 years of age and had metastatic ccRCC histologically confirmed, with the presence of measurable disease according to Response Evaluation Criteria in Solid Tumors v1.1. Details for SUVEGIL and TORAVA patient cohorts are described in supplementary Methods.

### Magnetic Resonance Imaging for in vivo mouse monitoring and mathematical modelling

Experiments were performed on a 7T Bruker BioSpec system equipped with a gradient coil of 660 mT.m-1 maximum strength and 110 µs rise time (see supplementary Methods for details).

### Additional statistical analysis

Additional statistical analysis (Besides R studio, see above) was performed using GraphPad Prism version 6.00 for Windows, GraphPad Software, La Jolla California USA, www.graphpad.com. For the statistical tests used, see figure legends.

## Acknowledgments

The Crispr/cas9 construct was gratefully provided by Michel Tremblay (Goodman Cancer Center, McGill, University). The authors would like to thank the lentiviral production plateform Vect’UB for EGFP lentivirus, the “service commun des animaleries”for animal production and housing, and the Bordeaux Imaging Center (BIC).

## Funding

This work was supported by grants from the PlanCancer («Systems Biology of Renal Cell Carcinoma using an Experimental RCC model»(C18005GS, SystemsRCC)), from the university Bordeaux (G2P project) and from the SIRIC BRIO project (Commucan) to AB. LC and AE were supported by post-doctoral fellowships from the Region “Nouvelle Aquitaine”.

## Author contributions

Study design: LSC, AB;experimental tasks:L.S.C., W.S, M.D., S.V., A.E., E.J.R., R.P., M-A.D.;data analysis: J.R., K.C. F.F., S.B., C.T., E.M., D.A., M.N.;writing of the manuscript: A.B., LSC, W.S. J.R., M.N.;technical support and conceptual advice: D.L.,, S.M. G.P.;provided clinical data and patient material: J-C.B., A.R., S.N., J-M.F.

## Competing interest

The authors declare no competing interest. Patents with regard to this article: 1)New gene signatures predicting survival time in patients suffering from renal cell carcinoma, EP19305300.6, March 13^th^, 2019;2) New Biomarkers and biotargets in renal cell carcinoma, EP19305252.9, March 5^th^, 2019.

## Data availability

Microarray gene expression data is available via Gene Expression Omnibus using the accession GSE142109 (Reviewer Access Token: upon request). DNA sequencing was deposited in ArrayExpress: http://www.ebi.ac.uk/arrayexpress/experiments/E-MTAB-8645, Access for reviewers: upon request. Methylation data is available via Gene Expression Omnibus using the accession GSE139338 (https://www.ncbi.nlm.nih.gov/geo/query/acc.cgi?acc=GSE139338;Reviewer Access Token: upon request).

## SUPPLEMENTARY MATERIALS

### Supplementary Methods

**For sub-cutaneous implantations of 786-O cells,** 5*10^6^ cells were injected into the flank of 5-week-old nude (nu/nu) female mice (Janvier, France). The tumor volume was determined with a caliper (v =L*l2*0.5). When the tumor reached 100 mm^3^, mice were treated once daily five times a week for 4 weeks by gavage with placebo (dextrose water vehicle) or sunitinib (40 mg/kg).

**Tissue dissociation and tumor cells purification.** For tumor cell purification, tissues were cut into small pieces with a scalpel and digested with Collagenase I and Collagenase II (Liberase TL, Roche, 05401020001) for 1 hour at 37°C. To further improve the dissociation, digested tissues were filtered in cell strainers (100µm, 70µm and 40µm) and seeded in complete medium, and incubated at 37°C, 5%CO_2_ in incubator. Cell cultures were checked daily and passaged as necessary. Tumor cell outgrowth and primary cell death resulted in tumor cell only cultures, verified by visualization of GFP using fluorescence microscopy When no GFP-negative cells could be visually detected, cell cultures were considered sufficiently pure. RENCA-GFP cells were collected for analysis or re-implanted into mice for the next *in-vivo* passage.

**Tissue processing.** For immunochemistry, tissue were fixed in paraformaldehyde 4%(PFA 4%, Santa Cruz Biotechnology, sc-281692) for 2 hours and then incubated for 72 hours in 30%sucrose. Tissues were frozen in OCT Compound (Tissue-Tek OCT compound, Sakura, 4583). Prior to embedding, lungs were inflated with 1mL of diluted OCT (1:1 PBS/OCT dilution). Frozen tissues were preserved at −80°C. For protein, DNA and RNA analysis, tissues were snap-frozen in liquid nitrogen and preserved at −80°C.

**Low-coverage genome sequencing.** Libraries were created using the KAPA LTP library preparation kit for Illumina platforms (Kapa Biosystems) following the manufacturers recommendations and sequenced on an Illumina HiSeq4000 sequencer in 51bp single-end mode. Raw reads were mapped to the mouse reference genome (mm10/GRCm38) using the Burrows-Wheeler Aligner *(Lin et al, 2019)*. On average, 6,882,693 reads were mapped per sample. PCR duplicates were removed using Picard (v1.43) resulting in, on average, 5,966,979 uniquely mapped reads per sample. These reads were further processed with the Bioconductor package QDNASeq.mm10. Problematic regions were excluded and read counts were corrected for mappability using LOESS regression. The number of reads was counted in bins of 50kb along the genome, log2-transformed and normalized by the median. These resulted in logR values per bin, that were subsequently segmented using the ASCAT algorithm *(Li &Durbin, 2009)*.These segmented values, as well as the individual logR values per bin, were used to plot the individual copy-number profile of each sample. Subclonal tumor fractions of the samples were estimated by ABSOLUTE *(Van Loo et al, 2010)*.

### Transcriptome data generation and analysis

From the log2 scale normalized data set, Principal Component Analysis was performed on series S0 and S2 as well as S0 and S5 (function prcomp of stats R package (v3.5.2) with the parameter center =T). Genes with the most important association where selected by keeping genes whose contribution is above the mean of all contributions for PC1 and/or PC2 for both S0 and S5 PCA analysis. This resulted in a set of 5140 genes. Expression values of these genes were further centered and reduced. For each of the series S0 through S5, a mean value was computed. We further built a heatmap of these mean values (pheatmap R package;v1.0.12).

| Group | S0 | S1 | S2 | S3 | S4 | S5 |
| --- | --- | --- | --- | --- | --- | --- |
| Parental cell line | 2 | NA | NA | NA | NA | NA |
| Kidney | NA | 2 | 2 | 3 | 5 | 5 |
| Tail | NA | 2 | 3 | 4 | 6 | 4 |
| Lung | NA | 1 | 5 | 3 | 4 | 3 |

**Methylomics data generation and analysis.** Whole Genome Bisulfite Sequencing (WGBS) was performed for 3 replicates of the S5 series for K, T, and L groups and 1 for the parental cell line at the GeT-PlaGe core facility, INRA Toulouse. WGBS libraries have been prepared according to Biooscientific’s protocol using the Biooscientific NEXTflex™Bisulfite Library Prep Kit for Illumina Sequencing. Briefly, DNA was fragmented by sonication, size selection was performed using Agencourt AMPure beads XP and adaptators were ligated to be sequenced. Then, bisulfite treatment was performed for 2.5 hours using the EZ DNA Methylation-Gold™Kit from Zymo Research, and 12 cycles of PCR were performed. Library quality was assessed using an Advanced Analytical Fragment Analyser and libraries were quantified by QPCR using the Kapa Library Quantification Kit. WGBS experiments have been performed on an Illumina HiSeq3000 using a paired end read length of 2×150 pb with the Illumina HiSeq3000 Reagent Kits. To determine conversion efficiency, fastq files were trimmed for adapters and low quality bases with Trim Galore (v0.4.4, calling cutadapt 1.3 *(Aken et al, 2017)* then mapped to the pUC19 reference genome (pUC19.fa) with Bismark (v0.13.0 *(Martin, 2011)*. Samtools (v0.1.19-44428cd *(Krueger &Andrews, 2011)* was used to remove duplicated reads. Then methylation calling was performed with Bismark_methylation_extractor. As methyled and non-methylated cytosine positions are known on the pUC19 reference genome, over and under-conversion could be assessed. Filtered fasq file were generated by CASAVA 2.17. Fastq files were aligned with Bismark (v0.17.1_dev *(Li et al, 2009)* against the GRCm38.p5 *mus musculus* genome (download from http://www.ensembl.org/, release 89 *(Krueger &Andrews, 2011)* with following parameters - N 0 and --maxins 800. Bismark use Bowtie 2 (v2.3.4.3 *(Aken et al, 2017)* and samtools (v1.9 *(Langmead et al, 2017)*. Incomplete bisulfite conversion filtering was done on Bismark BAM files in order to remove reads that exceed a certain threshold of methylated calls in non-CG context. Then, deduplication was applied (deduplicate_bismark) followed by extraction of methylated positions (bismark_methylation_extractor). Next analysis were done using R studio (R v3.5.2 *(R Foundation for Statistical Computing., 2018)*, R studio v1.1.463 *(Rstudio Team, 2016)*. We then considered CG positions with at least 10 reads of coverage resulting to 106303 CGs on chromosomes 1-19 and X. CGs were then annotated with RefSeq (download from UCSC website *(Li et al, 2009)* using the biomaRt R package (v2.38.0 *(Karolchik, 2004)* resulting in 6606 unique gene names. Principal Component Analysis (PCA) was performed on S0 and S5 for (methylation frequencies of all CGs (function prcomp of stats R package (v3.5.2) with the parameter center =T). CGs with the most important association were selected by keeping CGs whose contribution is up to 5 times the mean of all contributions for PC1 or PC2 of the PCA. The resulting 2063 CGs were localized in the gene body of unique 1427 genes used to perform enrichment analysis.

We further used the methylKit R package (v1.9.4) to calculate differential methylation at base resolution *(Durinck et al, 2009)*. After normalization of coverage values between samples by the median, reads covering both strands of a CpG dinucleotide were (merged. Fisher’s exact test was applied to identify differentially methylated cytosines (DMCs). Resulting DMCs in gene body were then confronted with genes that were differentially expressed between S0 and S5. TCGA methylation data sets (illuminaMethyl450_hg19_GPL16304_TCGAlegacy) of the KIRC cohort were download from XenaBrowser. Refseq (download for GRCh37 from UCSC website) was used to conserve only CGs that were in accord with the locus of the DMCs and only for genes that were in transcriptomics signatures. The resulting (5 CGs were located the intron 1 of IL34 and all had a positive correlation with IL34 expression. For each of these 5 CGs, we fit a Cox proportional hazard regression model based on overall survival (OS) (and disease-free survival (DFS). Two CGs had an HR >1 and a p-value of its log-rank test ≤0.01 for OS. As for transcriptomics data, the clinical relevance and prognostic performance of the 2 CGs were test against 1-CG random signatures using SigCheck R package. To separate samples into groups we computed a score corresponding to the mean value over all the methylation values for each sample ((scoreMethod=“High”in the sigCheck function). Patients were then split into 2 groups (low and high) to perform a log-rank test and compute associated HR (survival v2.44-1.1 and survminer v0.4.3 R packages). The Genome Browser screen shots were produced by using http://genome.ucsc.edu webtool (GRCm38 and GRCh37 *(Akalin et al, 2012)* and ensembl regulation data sets (mus_musculus.GRCm38.Regulatory_Build.regulatory_features) for GRCm38 and GeneHancer for GRCh37 *(Kent et al, 2002)*.

**Immunohistochemistry and immunofluorescence.** For frozen tissues, 10µm sections were performed with a cryostat (Leica CM1900).For paraffine tissues, 10µm sections were performed with a microtome. For frozen tissue immunofluorescence, sections were incubated 1 hour with a blocking buffer (5%BSA in PBS). Slides were incubated overnight with primary antibody (MMR R&D Systems, AF2535), and then with secondary fluorescent antibody (Interchim, FP547H) and DAPI (ThermoFisher Scientific, 62247). For paraffin tissues immunohistochemistry, slides were deparaffinized, re-hydrated and heated in Antigen Retrieval Solution pH6 (HIER Sodium Citrate Buffer, pH6;10mM Sodium Citrate, 0,05%Tween 20, pH 6,0). To block endogenous peroxidase activity, slices were treated with 0,3%hydrogen peroxide. After 1 hour of blocking in PBS 5%BSA, slides were incubated overnight with primary antibody (see table), and then incubated with biotinylated secondary antibody for 1h (see table). Secondary antibodies were HRP-conjugated using the “ABC”technique (Vectastain PK-6100) and then revealed with a peroxidase substrate kit (DAB, Vector Laboratories, SK-4100).

| Target | Antibody dilution | Reference | Secondary |
| --- | --- | --- | --- |
| Mouse MMR | 1/25 | R&D Systems, AF2535 | Interchim, FP-SA2110 |
| Human IL34 | 1/200 | Abcam, ab101443 | Dako, E0433 |

Bright-field microscopy was performed with a Nikon Eclipse E4000 microscope (Nikon) and NIS-Elements F 3.2 software (Nikon). Fluorescent microscopy was performed with a Nikon Eclipse i90 microscope (Nikon) and NIS-Elements AR 4.30 software (Nikon). A slide scanner (Hamamatsu, Nanozoomer 2.0HT) from the Bordeaux Imaging Center was used for whole slide imaging using NDP.scan software (Hamamatsu). Analysis were performed using Fiji. Cell counting was performed with “Cell Counter”plugin (Kurt De Vos).

**Western Blot Analysis.** Cells were washed with PBS and dissolved in lysis buffer (50mM Tris-HCl pH 7,4, 150mM NaCl, 1%TritonX-100, 0,1%SDS, 0,1%sodium deoxycholate) supplemented with protease and phosphatase inhibitors cocktails (Roche, Complete Protease Inhibitor Cocktail;PhosSTOP). For tissues, pieces of approximately 5mm^3^ were dissolved in lysis buffer. Protein concentration was quantified by Bradford assay (Euromedex). Protein lysates were diluted in Laemmli buffer (62,5mM Tris-HCl pH 6,8, 10%glycerol, 2,5%SDS, 2,5%β-mercaptoethanol), boiled for 5 minutes at 95°C and resolved by SDS-PAGE. Protein were then transferred onto a nitrocellulose membrane (BioTrace, 66485). Membranes were incubated with blocking buffer (see table) for 1 hour, and then probed with primary antibody (see table).

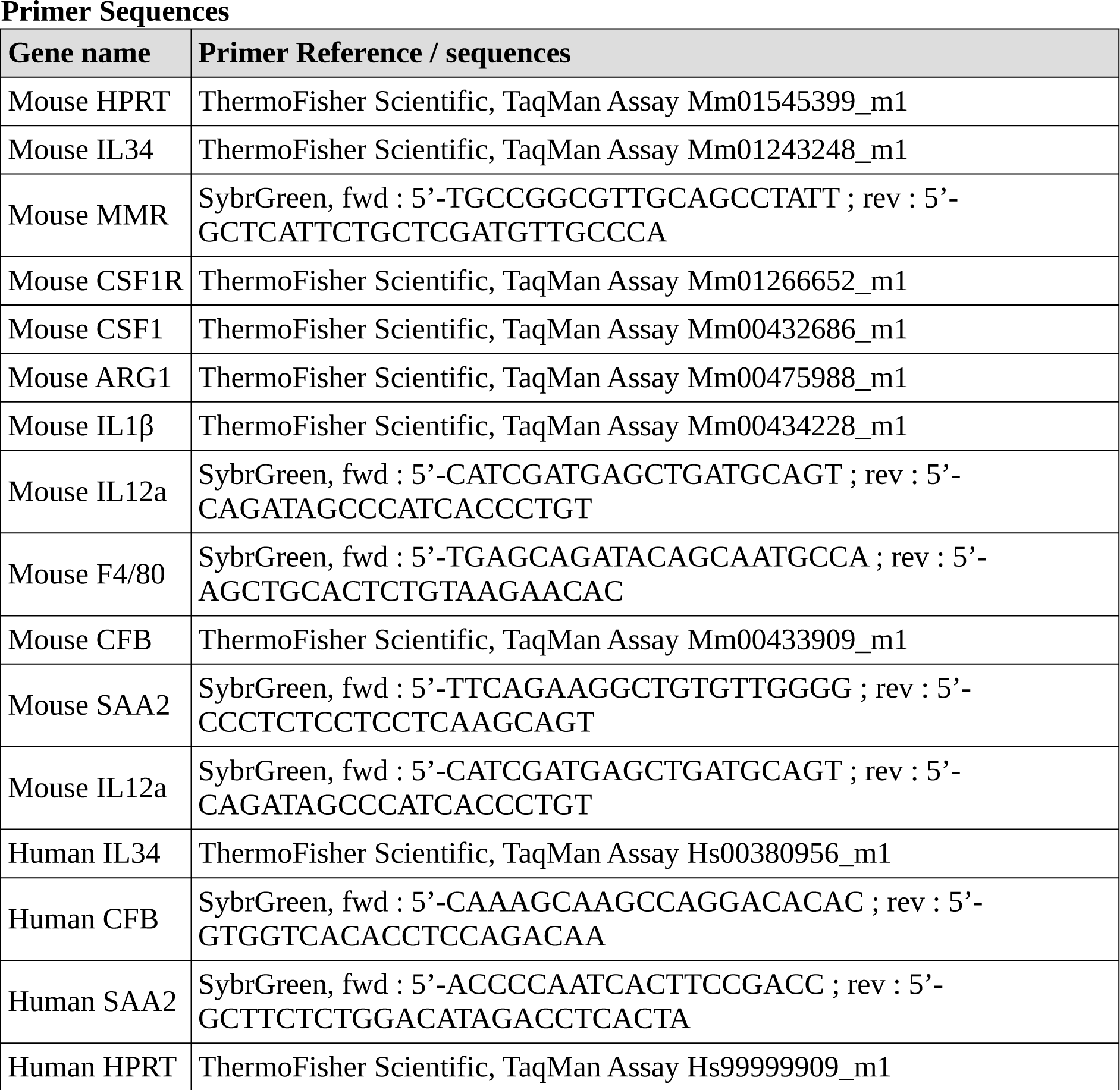

### Generation of GFP, IL34, overexpressing cells and crispr/cas9 induced IL34 knock-down cells

EGFP expressing lentivirus (pRRLsin-MND-eGFP-WPRE) was obtained from the vectorology plateform of the University of Bordeaux (Vect’UB).

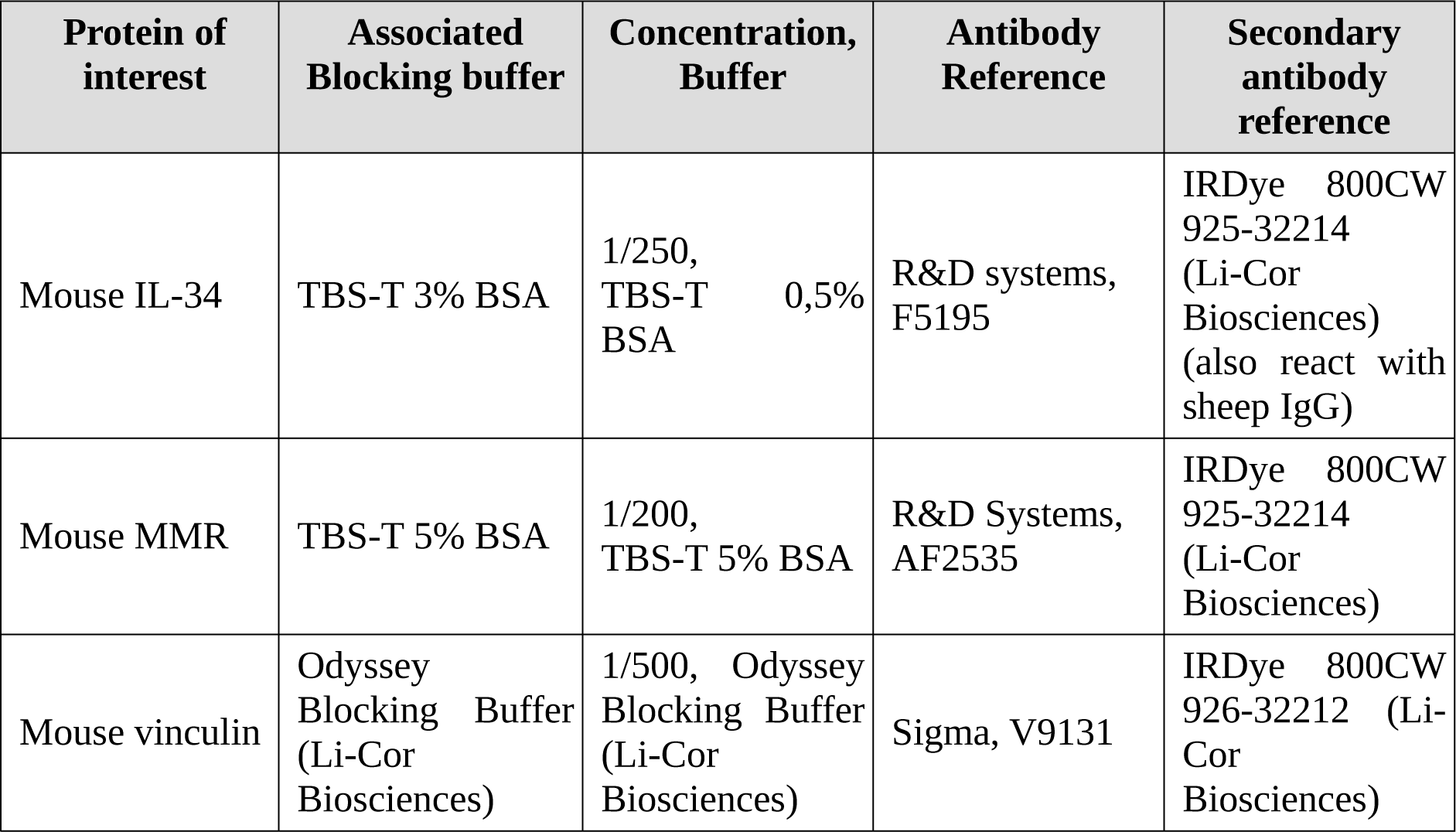

The IL34 lentiviral vector was obtained the following way. Total RNA was extracted from Renca cell and reverse-transcribed into cDNA using the high-capacity cDNA reverse transcription kit (Applied Biosystems). Mouse IL34 variant 1 cDNA were then amplified by PCR using specific primers containing attB site: fwd: 5’-GGGG-ACA-AGT-TTG-TAC-AAA-AAA-GCA-GGC-TTC-ATG-CCC-TGG-GGA-CTC-GCC-TGG-CTA-3’;rev: 5’-GGGG-AC-CAC-TTT-GTA-CAA-GAA-AGC-TGG-GTC-TCA-GGG-CAA-CGA-GCC-ATG-GCT-TGA-3’. attB-PCR product was cloned into pDONR221 (ThermoFisher, #12536017) by BP gateway cloning and then transferred into pLenti CMV/TO Puro DEST (Addgene #17293) by LR gateway cloning to give pLenti CMV/TO-mIL34v1-Puro lentiviral vector. For the IL34 lentiviral cripr/cas9 construct, lentiCRISPRv2 plasmid containing spCas9 cassette and expressing guide RNA targeted LacZ or mouse IL34 exon 1 were kindly provided by Michel Tremblay (McGill University) with the following targeted sequence: mIL34 ex1a: 5’-gatcctacttgacgtggctttgg-3’and mIL34 ex1c: 5’-gaccttacaggctaccttcgggg-3’. Viral particles were produced by calcium phosphate co-transfection of HEK293T cells with the packaging plasmids pVSVg (Addgene 8454), psPAX2 (Addgene 12260) and a transfer plasmid (e.g. EGFP- or mIL34v1-plasmid or pLentiCRISPRv2-LacZ, -mIL34_1a, - mIL34_1c). Renca cells were infected with lentiviruses at a multiplicity of 25 infectious units per cell and selected for 4 days in media supplemented with 1µg/ml puromycin.

**Additional details for human patient samples.** Patients had not received previous systemic therapy for RCC and were eligible for sunitinib or bevacizumab treatment in the first-line setting. Patients were ineligible if they had symptomatic or uncontrolled brain metastases, an estimated lifetime less than 3 months, uncontrolled hypertension or clinically significant cardiovascular events (heart failure, prolongation of the QT interval), history of other primary cancer. All patients gave written informed consent. Tumors were assessed at baseline and then every 12 weeks by thoracic, abdominal, pelvic and bone CT scans. Brain CT scans were performed in case of symptoms. This cohort includes patients from the SUVEGIL (24 patients) and TORAVA (35 patients) trials. The SUVEGIL trial (clinicaltrial.gov, <u>NCT00943839</u>) was a multi-center prospective single-arm study. The goal of the trial is to determine whether a link exists between the effectiveness of therapy with sunitinib malate and development of blood biomarkers in patients with kidney cancer. Patients received oral sunitinib (50 mg per day) once daily for 4 weeks (on days 1 to 28), followed by 2 weeks without treatment. Courses repeat every 6 weeks in the absence of disease progression or unacceptable toxicity. The TORAVA trial (clinicaltrial.gov, <u>NCT00619268</u>) was a randomized prospective study. Patient characteristics and results have been previously described *(Bult et al, 2019)*. Briefly, patients aged 18 years or older with untreated metastatic ccRCC were randomly assigned (2: 1: 1) to receive the combination of bevacizumab (10 mg kg^−1^ iv every 2 weeks) and temsirolimus (25 mg iv weekly) IFN-α(9 mIU i.v. three times per week), or one of the standard treatments: sunitinib (50 mg per day orally for 4 weeks followed by 2 weeks off) *(Négrier et al, 2011)*. These studies were approved by the ethic committee at each participating center and run in agreement with the International Conference on Harmonization of Good Clinical Practice Guideline. Blood samples were collected during the inclusion visit (baseline).

**Magnetic Resonance Imaging for in vivo mouse monitoring.** A volume resonator operating in quadrature mode was used for excitation (75.4mm inner diameter, 70mm active length), and a proton phased array (RAPID Biomedical GmbH) was used for signal reception (4 elements of 30 mm long around an elliptic cylinder housing: 19 ×25.5 mm). A Self-Gating balanced Steady State Free Precession sequence (SG-bSSP *(Ribot et al, 2015)*) was employed to image the mouse lungs in order to count the pulmonary metastases. The following parameters were used: Field-Of-View (FOV): 25×22×20mm, spatial resolution: 195×172×156um, TE/TR=2/4ms, reception bandwidth (BW): 100kHz, flip angle: 30°, 4 repetitions, 4 phase offsets, total acquisition time: 22min. Thereafter, the motion correction technique was used as already explained in Ribot EJ et al *(Ribot et al, 2015)*. A standard bSSFP sequence was also applied to image the mouse abdomen and measure the primary tumor volumes. The following parameters were used: FOV: 25×22×20mm, spatial resolution: 195×172×156um, TE/TR=2/4ms, BW: 75kHz, flip angle: 30°, 4 averages, 4 phase offsets, total acquisition time: 22min.

**Mathematical modeling methods.** The data consisted of longitudinal observations of total volume of the kidney and the primary tumor (PT) coupled with lung metastases counts, for two groups of six mice, the control group LacZ and the IL34 KO CRISPR group. These data were obtained from MRI images of subcapsular implanted mice. The volumes of the mice kidneys were measured and were approximately equal to 187 mm^3^, enabling then to deduce the PT volume for each mouse. We considered a model that included two ordinary differential equations: one for the primary tumor growth, and the other for metastatic invasion (counts of metastases) as a result of dissemination from the PT. We denoted by *V*_*p*_ and *N* the volume of the PT and the number of lung metastases, respectively. The initial time *t*^0^=0 is the time of tumor cells implantation. The initial volume of the primary tumor *V*_*p*_^0^ is set at the volume converted from the number of injected cells (using the conversion rule 1 mm^3^ ≈10^6^ cells *(Spratt et al, 1995)*, thus *V_p_*0=10^5^ cells here. The model writes

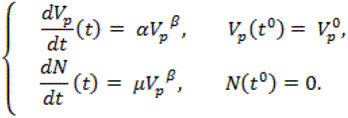

The first equation describes the growth kinetics of the PT with a growth rate that depends on the volume of the PT. It is assumed to follow a power law model *(Benzekry et al, 2014)* with parameters *α* (proliferation rate) and *β* (fractal dimension of the PT vasculature). The second equation features the metastatic development through a PT volume-dependent rate, defined by the parameters *μ* and *β*. The parameter *μ* is the per cell probability of successful emission of metastases from the primary tumor to the lungs. We refer to this parameter as the intrinsic dissemination coefficient. The same fractal dimension *β* is used following the assumption that cells able to leave are cells close to the vasculature *(Iwata et al, 2000)*. In order to calibrate the mathematical models with the data, we used a statistical population approach by means of the non-linear mixed effects formalism *(Lavielle, 2014)*. We assumed log-normal distributions for the parameters (*α*^*μ*^,*μ*^*μ*^) with *μ* the animal index, with group effect considered as a categorical on *α* and *μ*. Parameter *β* was found to exhibit small variability and no random effects were considered for this parameter.

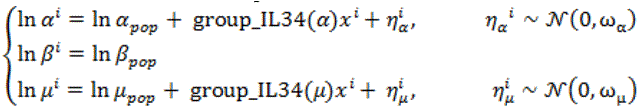

where x^*i*^=1 if animal *μ* is in the IL34 KO group and 0 if not.We assumed a proportional error model for the PT volume observations and a constant error model for the metastases counts. To compute the parameters of interest, we maximized the likelihood of the pooled data using the stochastic approximation expectation maximization (SAEM) algorithm implemented within the Monolix 2019R1 software (Lixoft SAS).

## Supplementary Figures

**Fig. S1.**
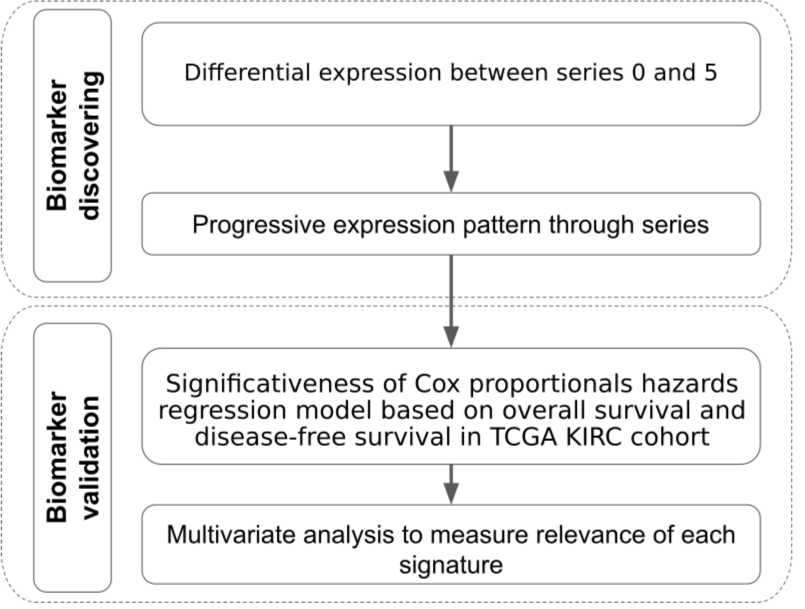
**The figure depicts the general strategy used to determine clinically-relevant signatures from the transcriptomic data.**

**Fig. S2.**
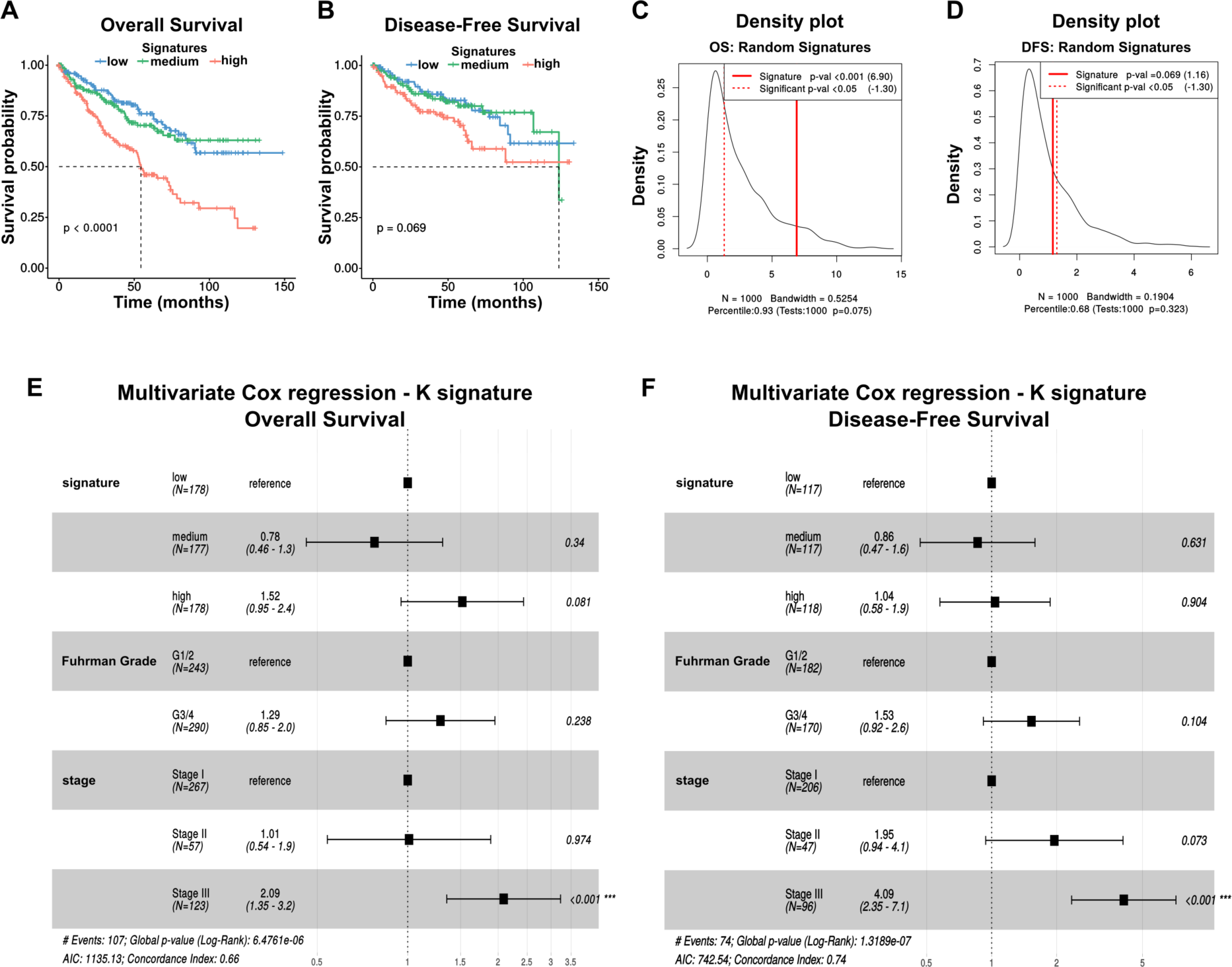
Validation of K signature in the KIRC TCGA cohort. (**A** and **B**) Kaplan-Meier for overall (OS) and disease-free (DFS) survival analysis stratified in 3 groups of equivalent size. Signature “low”: patient group with low score;signature “medium”: patient group with a medium score;signature “high”: patient group with high score. (**C** and **D**) Density plot of p values (log-rank test) from 1000 random signatures of equal size for OS and DFS. p =empirical p-value. (**E** and **F**) Forest plot of multivariate Cox proportional hazards model for the Lung signature in the KIRC cohort adjusted for clinical variables (Fuhrman grade and TNM stage).

**Fig. S3.**
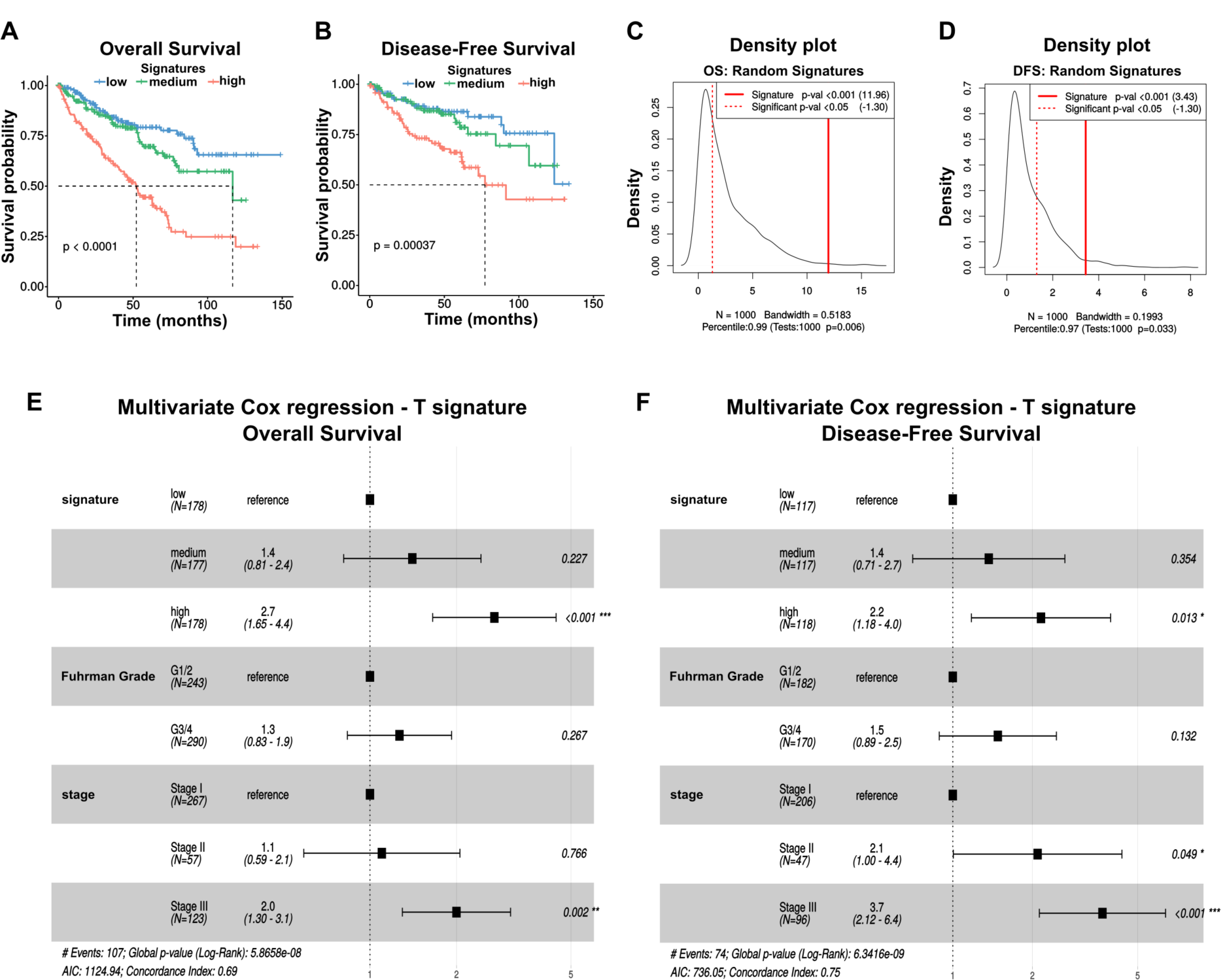
Validation of T signature in the KIRC TCGA cohort. (**A** and **B**) Kaplan-Meier for overall (OS) and disease-free (DFS) survival analysis stratified in 3 groups of equivalent size. Signature “low”: patient group with low score;signature “medium”: patient group with a medium score;signature “high”: patient group with high score. (**C** and **D**) Density plot of p values (log-rank test) from 1000 random signatures of equal size for OS and DFS. p =empirical p-value. (**E** and **F**) Forest plot of multivariate Cox proportional hazards model for the Lung signature in the KIRC cohort adjusted for clinical variables (Fuhrman grade and TNM stage).

**Fig. S4.**
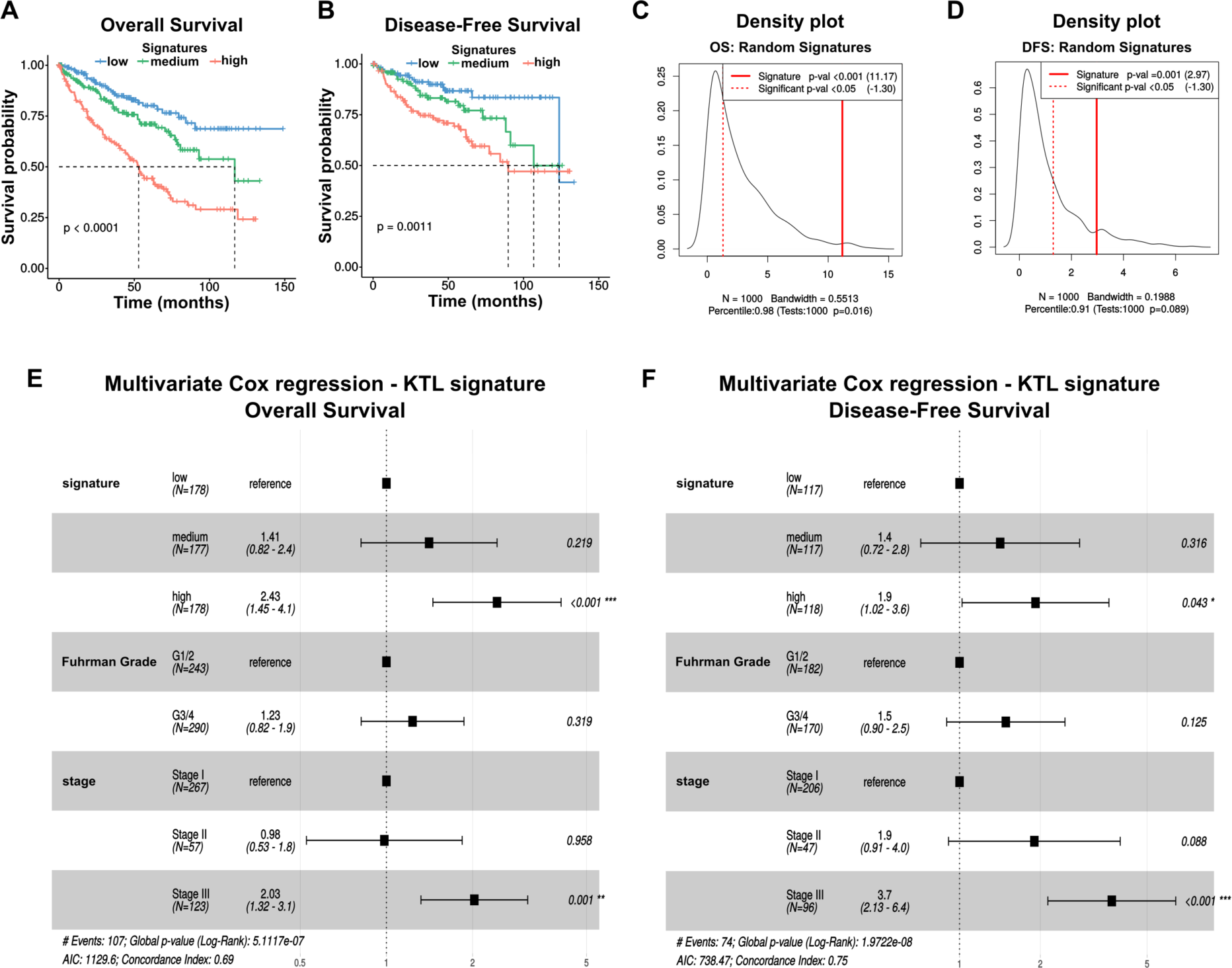
Validation of KTL signature in the KIRC TCGA cohort. (**A** and **B**) Kaplan-Meier for overall (OS) and disease-free (DFS) survival analysis stratified in 3 groups of equivalent size. Signature “low”: patient group with low score;signature “medium”: patient group with a medium score;signature “high”: patient group with high score. (**C** and **D**) Density plot of p values (log-rank test) from 1000 random signatures of equal size for OS and DFS. p =empirical p-value. (**E** and **F**) Forest plot of multivariate Cox proportional hazards model for the Lung signature in the KIRC cohort adjusted for clinical variables (Fuhrman grade and TNM stage).

**Fig. S5.**
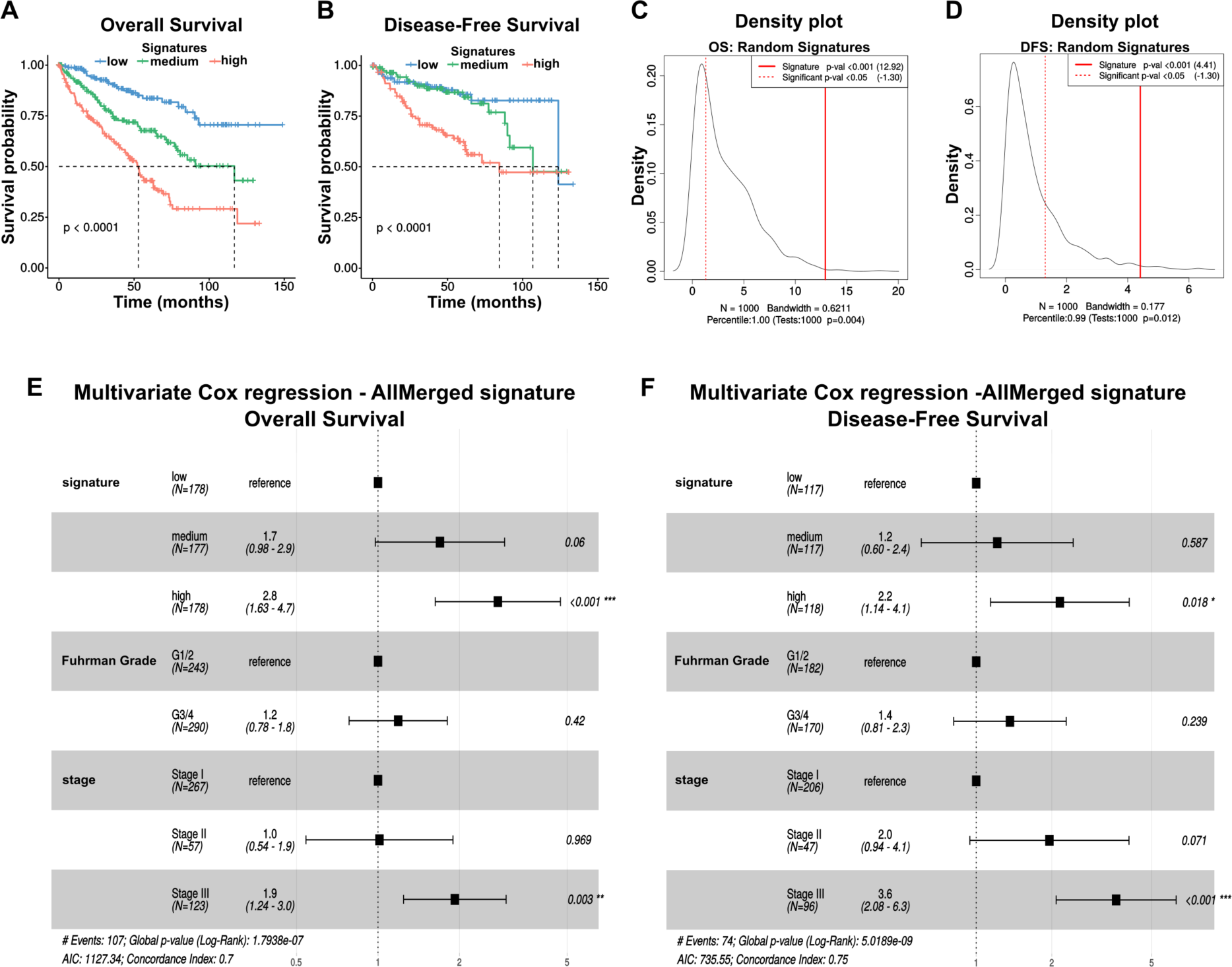
Validation of AllMerged signature in the KIRC TCGA cohort. (**A** and **B**) Kaplan-Meier for overall (OS) and disease-free (DFS) survival analysis stratified in 3 groups of equivalent size. Signature “low”: patient group with low score;signature “medium”: patient group with a medium score;signature “high”: patient group with high score. (**C** and **D**) Density plot of p values (log-rank test) from 1000 random signatures of equal size for OS and DFS. p =empirical p-value. (**E** and **F**) Forest plot of multivariate Cox proportional hazards model for the Lung signature in the KIRC cohort adjusted for clinical variables (Fuhrman grade and TNM stage).

**Fig. S6.**
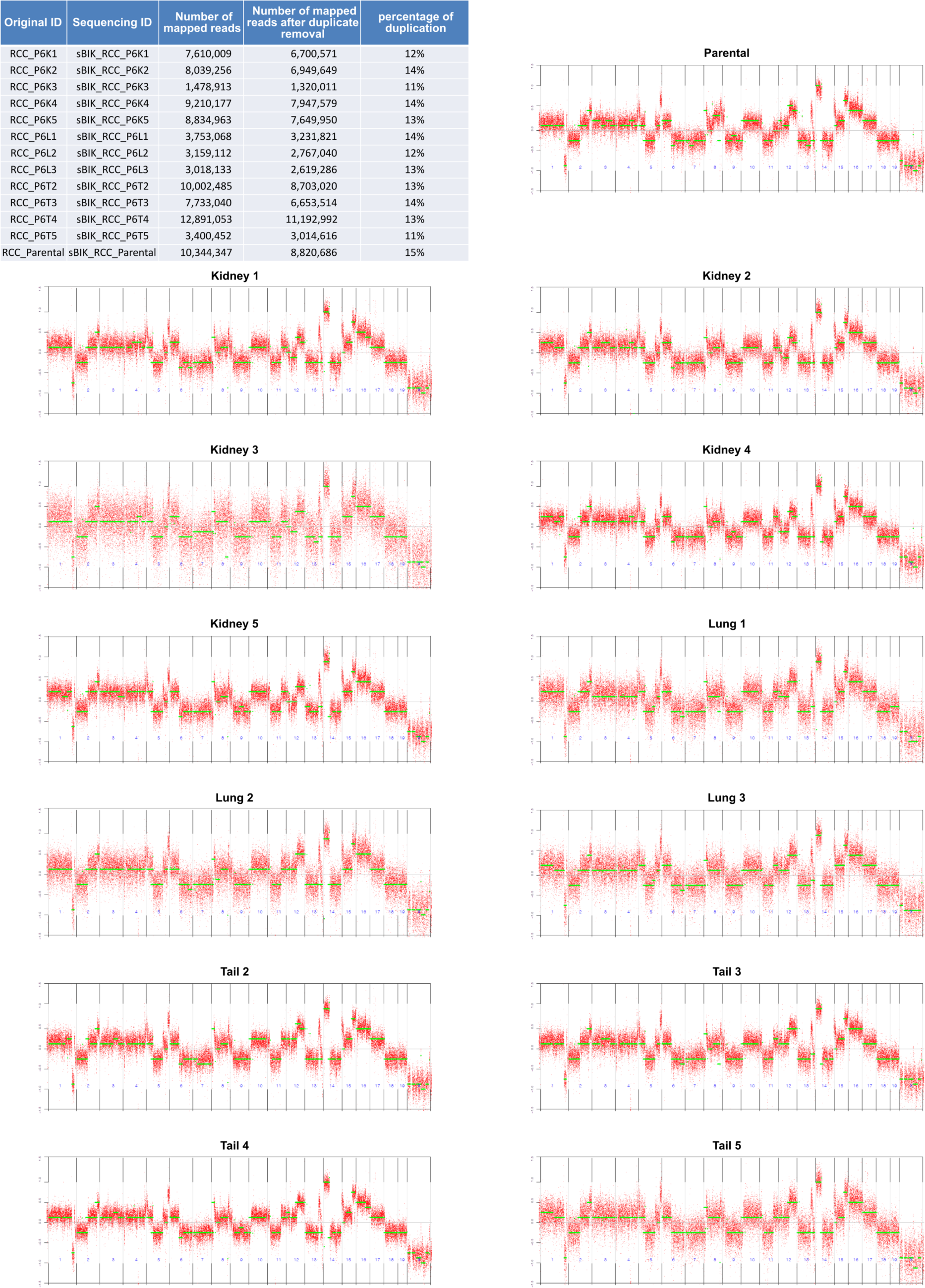
Low-coverage shallow DNA sequencing of the S5 series versus parental cells. 5 Kidney, 3 Lungs and 4 tail samples were sequenced. There is no significant copy number variation in these different groups.

**Fig. S7.**
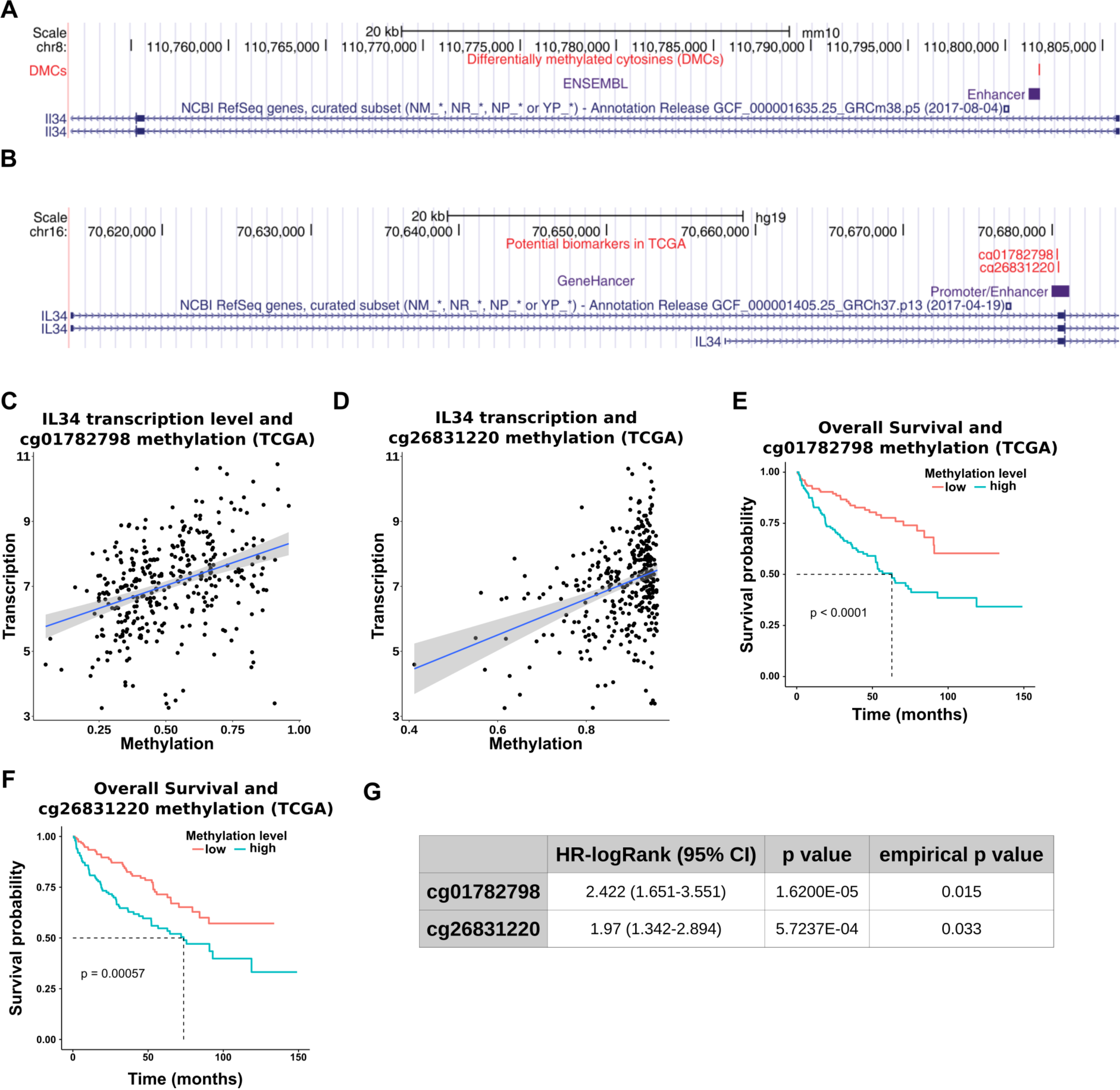
IL34 CpG methylation and clinical relevance. (**A**) Locus of the identified DMC in intron 1 of IL34 (GRCm38) overlapping an enhancer (ENSEMBL) (**B**) Locus of the CpGs cg01782798 and cg26831220 in the intron 1 of IL34 (GRCh37) overlapping an enhancer (GeneHancer);(**C** and **D)** Scatterplot comparing the IL34 transcription and methylation level of the CpGs;(**E** and **F**) Kaplan-Meier plots for overall survival stratified in 2 groups (TCGA KIRC cohort). low: patient group with low methylation level;high: patient group with high methylation level. (**G**) Summary of results. HR (95%CI): Hazard Ratio with 95%Confidence Interval;p value: log-rank test p-value;empirical p value: prognostic performance of the 2 CGs were test against 1-CG random signatures.

**Fig. S8.**
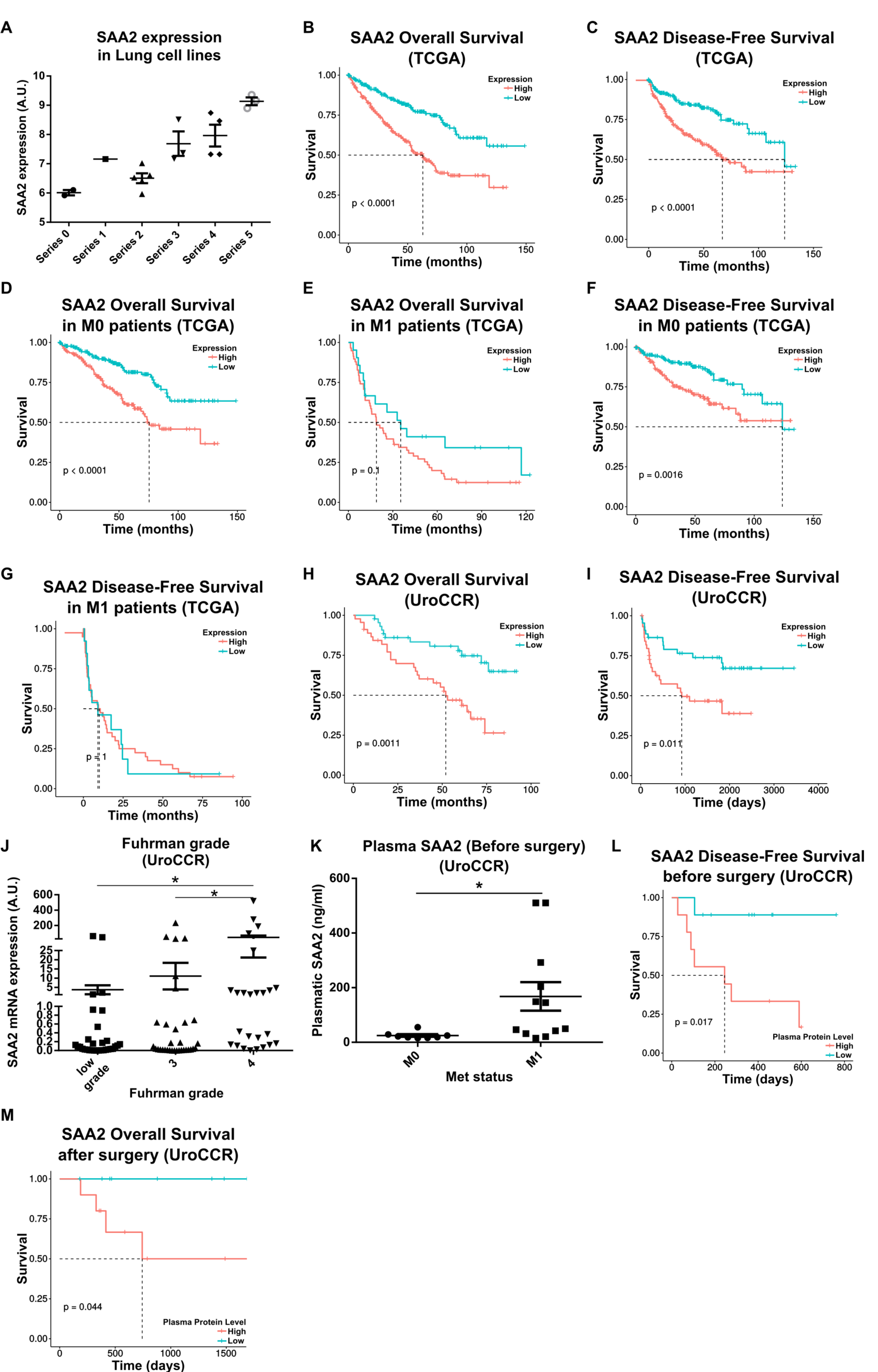
SAA2 and clinical relevance. (**A**) SAA2 expression in Lung cell lines with increasing passages compared to parental cells (Series 0). graph shows means ±SEM. (**B**) Overall Survival (OS) of patients stratified according to SAA2 expression (TCGA KIRC Cohort). (HR(log-rank)=2.472 (1.835-3.329)) (**C**) Disease Free Survival (DFS) of patients stratified according to SAA2 expression (TCGA KIRC Cohort)(HR(log-rank)=3.562 (2.278-5.57). (**D** to **G**) OS and DFS of patients stratified according to SAA2 expression (TCGA KIRC Cohort) and their metastatic status(OS: M0 HR(log-rank)=2.334 (1.591-3.423);M1[NS] HR(log-rank)=1.62 (0.955-2.749) –DFS: M0 HR(log-rank)=3.511 (1.851-6.658);M1[NS] HR(log-rank)=2.055 (1.038-4.07)). (**H**) Overall Survival (OS) of patients stratified according to SAA2 expression in the UroCCR cohort (HR(log-rank)=2.901 (1.526-5.517)). (**I**) Disease Free Survival (DFS) of patients stratified according to SAA2 expression in the UroCCR cohort (HR(log-rank)=2.342 (1.211-4.529)). (**J**) Correlation between SAA2 expression in patients and the Fuhrman grade (UroCCR cohort). (**K**) Plasma SAA2 levels in non-metastatic (M0) and metastatic (M1) patients at diagnosis in the UroCCR cohort. (**L**) DFS and correlation with pre-surgery SAA2 plasma levels (UroCCR cohort)(HR(log-rank)=8.191 (2.04-32.891)). (**M**) OS and correlation with post-surgical SAA2 plasma levels (UroCCR cohort)(HR(log-rank)=Inf (Inf-Inf)[not calculable]).

**Fig. S9.**
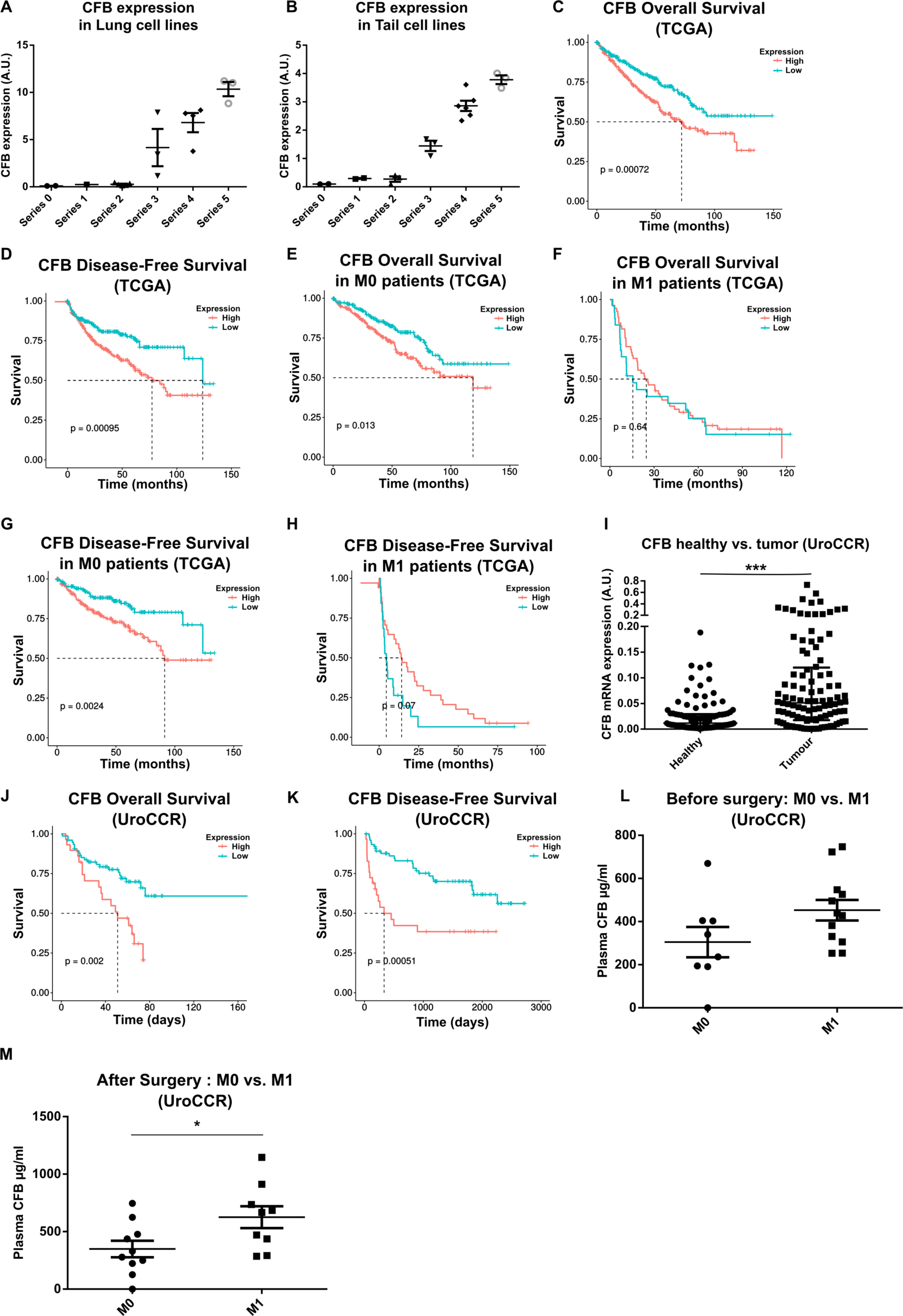
CFB and Clinical Relevance. (**A)** CFB expression in Lung and (**B**) Tail cell lines with increasing passage compared to parental cells (Series 0). graph shows means ±SEM. (**C**) Correlation of CFB to OS in patients from the TCGA KIRC Cohort.(HR(log-rank)=1.684 (1.251-2.266)) (**D**) Correlation of CFB to DFS in patients from the TCGA KIRC Cohort (HR(log-rank)=1.935 (1.238-3.025)). (**E** to **H**) Correlation of CFB to OS and DFS in metastatic and non-metastatic patients patients from the TCGA KIRC Cohort (OS: M0 HR(log-rank)=1.616 (1.105-2.363);M1[NS] HR(log-rank)=0.883 (0.514-1.518) –DFS: M0 HR(log-rank)=2.605 (1.379-4.921);M1[NS] HR(log-rank)=0.771 (0.388-1.532)). (**I**) CFB expression in tumor and healthy tissues from UroCCR cohort. (**J**) Correlation of CFB to OS in patients from the UroCCR cohort (HR(log-rank)=2.556 (1.24-5.267)). (**K**) Correlation of CFB to DFS in patients from the UroCCR cohort.(HR(log-rank)=2.846 (1.323-6.123)). (**L** and **M**) CFB plasma level in patients from the UroCCR cohort before and after surgery.

**Fig. S10.**
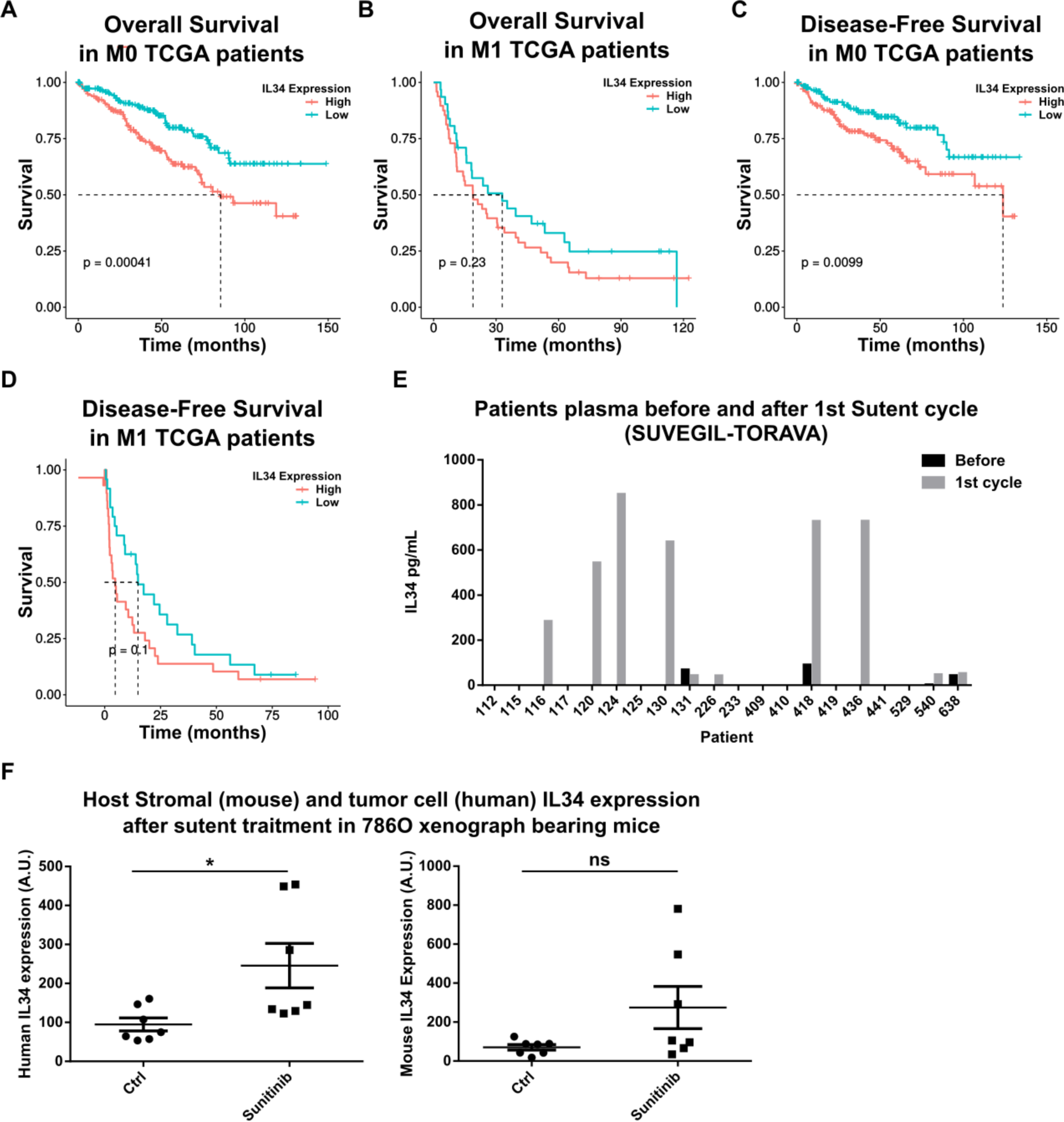
IL34 and clinical correlations. (**A** to **D**) OS and DFS of patients stratified according to IL34 expression (TCGA KIRC Cohort) and their metastatic status (OS: M0 HR(log-rank)=1.985 (1.359-2.9);M1[NS] HR(log-rank)=1.368 (0.834-2.242) –DFS: M0 HR(log-rank)=3.542 (1.874-6.694);M1[NS] HR(log-rank)=1.38 (0.731-2.606)). (**E**) IL34 plasmatic levels in patients before and after sunitinib treatment (SUVEGIL-TORAVA cohorts) (**F**) IL34 mRNA expression in mice implanted with 786-O cells and treated with sunitinib. Specific primers were used for human (left panel) and mouse (right panel) IL34. Results presented as mean±SEM. *p<0.05. ns:Non significant. Unpaired t test.

**Fig. S11.**
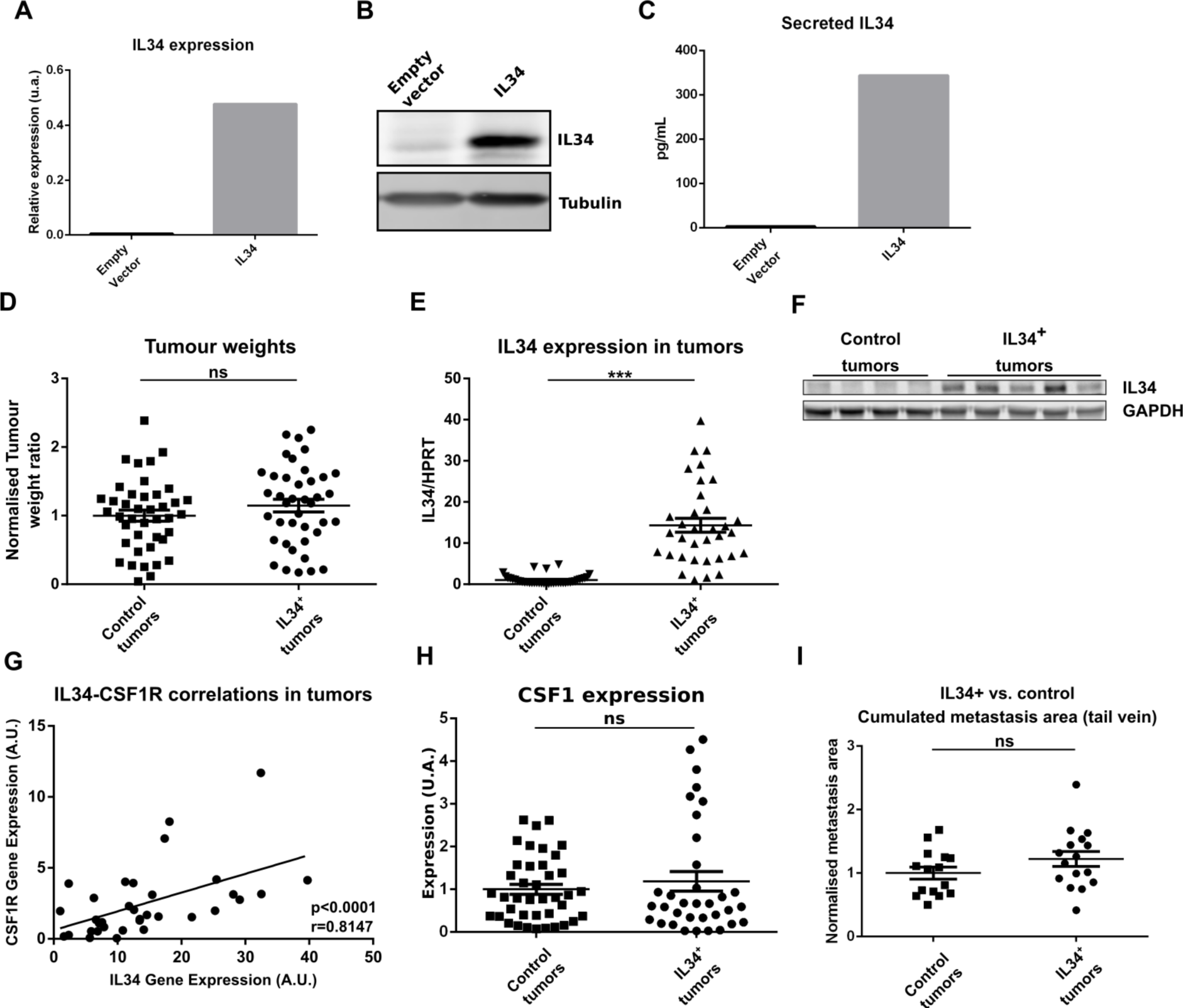
IL34 overexpression in renal carcinoma cells. (**A** and **B**) Validation of IL34 expression by qPCR and western-blot in RENCA cells transduced with lentiviral vector (pLenti CMV/TO-mIL34v1-Puro) encoding for mouse IL34. (**C**) IL34 ELISA assay from supernatants of transduced cells. (**D**) Tumor weights of IL34 overexpressing and control tumors. (**E** and **F**) Validation of IL34 expression in IL34 overexpressing tumors by qPCR (T-test. ***p<0.001) and Western Blot. (**G**) Correlation between IL34 mRNA and CSF1R mRNA (global macrophage marker). And Spearman correlation test. (**H**) CSF1 mRNA expression in IL34 overexpressing and control tumors. (**I**) Quantification of the metastasis area in lung sections after tail vein injection of IL34 overexpressing or control cells. T-test (ns).

**Fig. S12.**
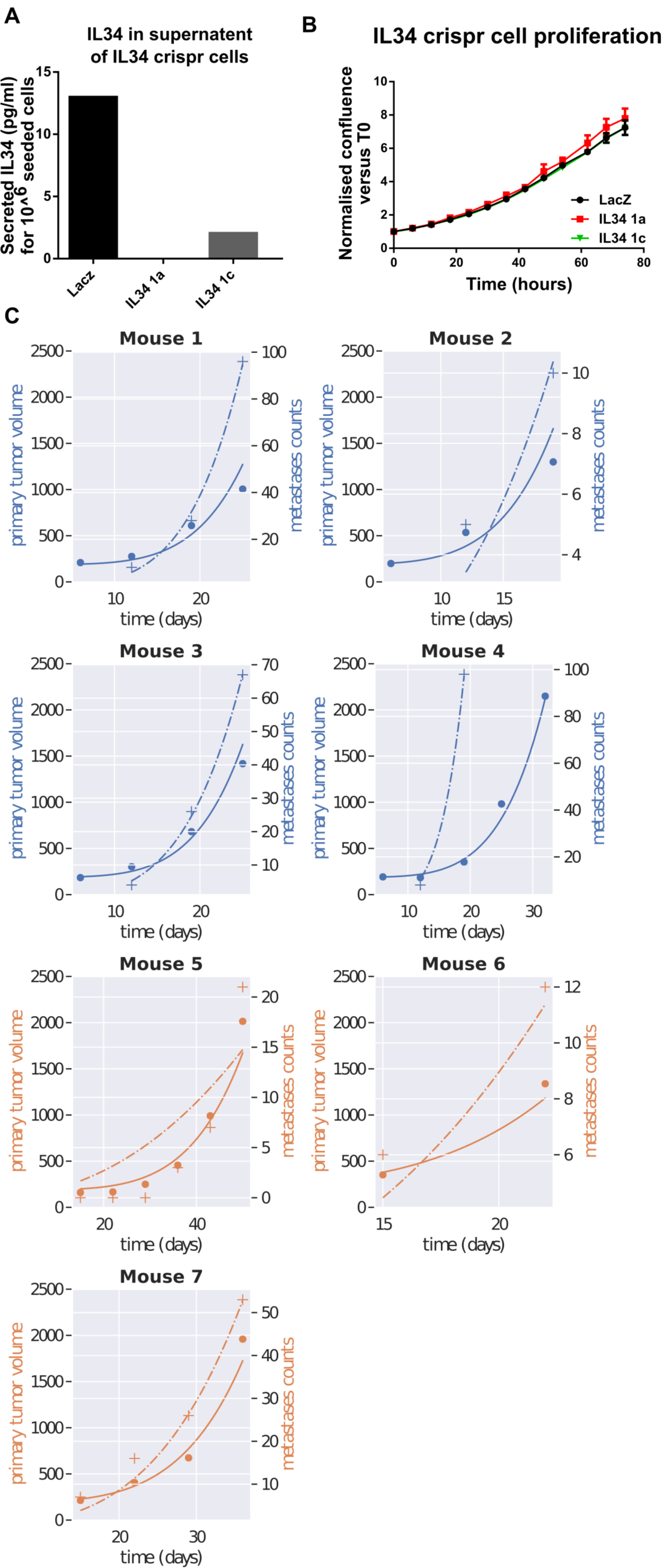
Validation of IL34 knock-out and mathematical modeling. (**A**) validation of IL34 knock-down by ELISA in conditioned media of RENCA cells. (**B**) Proliferation assay with crispr IL34-ko (IL341a, IL341b) and control cells (LacZ) with no significant differences (Dunnett’s multiple comparison tests on all timepoints). (**C**) Mathematical modeling of primary tumor growth kinetics for 7 different animals.

**Table S1.**
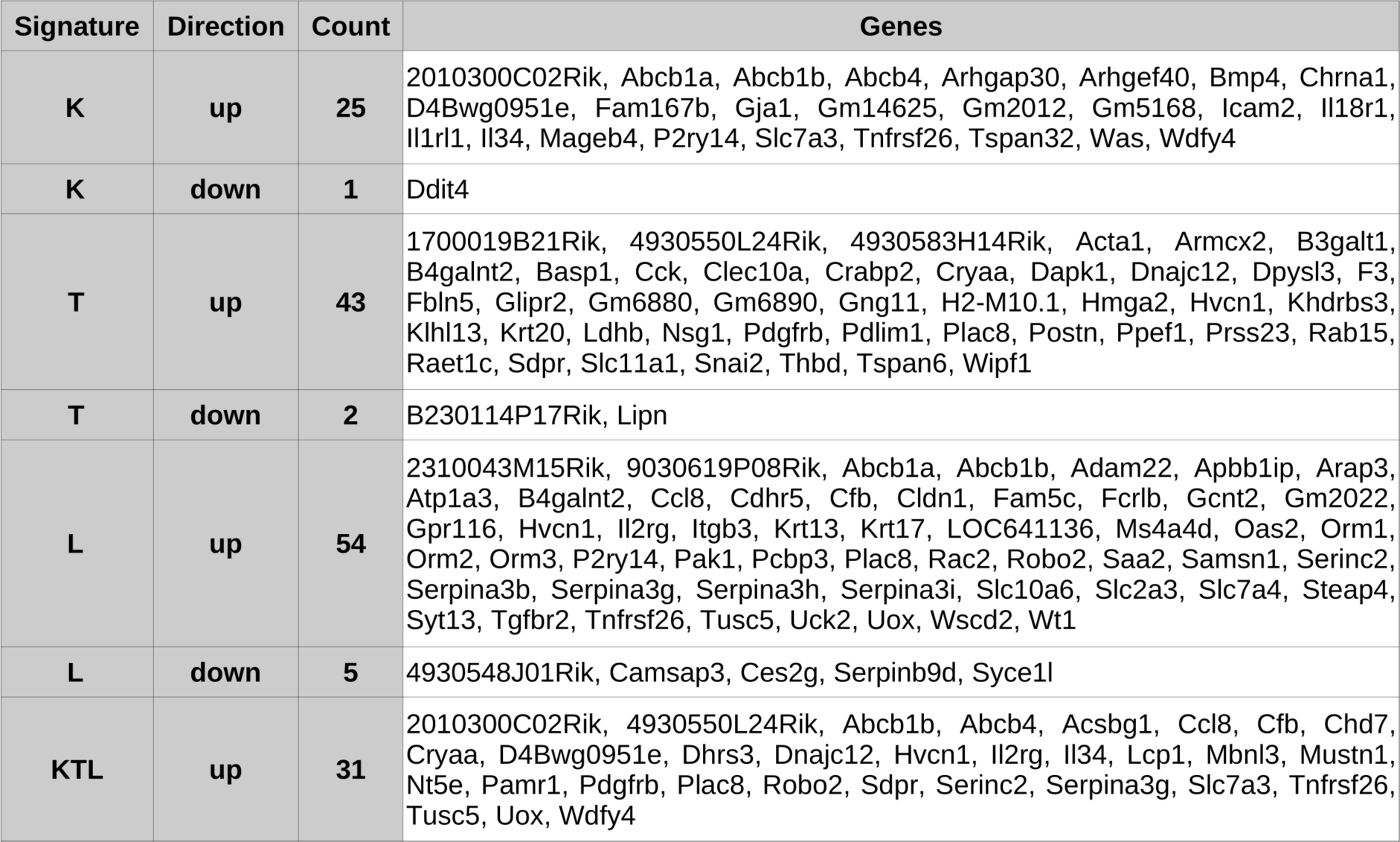
Summary table of up and downregulated genes from the first biomarker discovery step.

**Table S2.** **List of genes present for each signature with details of the predictive value.**

| Signature | Gene | log2(FC) | Human Gene Name | OS |  |  | DFS |  |  |
| --- | --- | --- | --- | --- | --- | --- | --- | --- | --- |
|  |  |  |  | HR (95% CI) | pvalue | FDR | HR (95% CI) | pvalue | FDR |
| K | Il34 | 4.021E+00 | IL34 | 1.287 (1.137-1.458) | 5.197E-05 | 5.544E-04 | 1.362 (1.176-1.578) | 3.216E-05 | 5.544E-04 |
| K | Mageb4 | 4.614E+00 | MAGEB2 | 1.239 (1.141-1.345) | 4.837E-05 | 5.544E-04 | 1.213 (1.096-1.342) | 2.568E-03 | 1.027E-02 |
| K | Tspan32 | 2.870E+00 | TSPAN32 | 1.261 (1.11-1.432) | 3.347E-04 | 2.142E-03 | 1.04 (0.895-1.208) | 6.078E-01 | 8.104E-01 |
| K | Was | 2.082E+00 | WAS | 1.337 (1.144-1.562) | 1.719E-04 | 1.375E-03 | 1.334 (1.105-1.611) | 1.980E-03 | 9.052E-03 |
| T | 4930550L24Rik | 3.870E+00 | MAGEC2 | 1.282 (1.182-1.392) | 3.038E-06 | 2.891E-05 | 1.292 (1.146-1.457) | 8.876E-04 | 2.515E-03 |
| T | 4930550L24Rik | 3.870E+00 | MAGEC3 | 1.216 (1.119-1.321) | 5.181E-05 | 2.202E-04 | 1.17 (1.05-1.304) | 1.096E-02 | 2.405E-02 |
| T | Basp1 | 2.164E+00 | BASP1 | 1.409 (1.264-1.571) | 1.638E-09 | 5.568E-08 | 1.375 (1.203-1.573) | 5.520E-06 | 3.412E-05 |
| T | Crabp2 | 4.038E+00 | CRABP2 | 1.196 (1.131-1.266) | 4.139E-09 | 7.940E-08 | 1.133 (1.057-1.216) | 7.903E-04 | 2.337E-03 |
| T | Dnajc12 | 3.106E+00 | DNAJC12 | 1.246 (1.139-1.363) | 4.973E-06 | 3.381E-05 | 1.24 (1.116-1.377) | 1.441E-04 | 5.443E-04 |
| T | Dpysl3 | 4.673E+00 | DPYSL3 | 1.341 (1.185-1.517) | 4.665E-06 | 3.381E-05 | 1.498 (1.304-1.721) | 3.628E-08 | 4.935E-07 |
| T | F3 | 4.971E+00 | F3 | 1.209 (1.115-1.312) | 1.034E-05 | 5.861E-05 | 1.22 (1.097-1.357) | 3.947E-04 | 1.220E-03 |
| T | Glipr2 | 2.374E+00 | GLIPR2 | 1.562 (1.274-1.917) | 1.374E-05 | 7.185E-05 | 1.465 (1.156-1.856) | 1.321E-03 | 3.593E-03 |
| T | Hmga2 | 2.645E+00 | HMGA2 | 1.199 (1.137-1.265) | 4.670E-09 | 7.940E-08 | 1.164 (1.084-1.249) | 1.754E-04 | 6.276E-04 |
| T | Ppef1 | 4.674E+00 | PPEF1 | 1.278 (1.159-1.41) | 1.743E-06 | 1.976E-05 | 1.307 (1.173-1.457) | 3.402E-06 | 2.891E-05 |
| T | Rab15 | 4.222E+00 | RAB15 | 1.388 (1.186-1.626) | 4.876E-05 | 2.202E-04 | 1.196 (0.992-1.441) | 6.103E-02 | 1.092E-01 |
| T | Slc11a1 | 3.492E+00 | SLC11A1 | 1.469 (1.3-1.661) | 1.143E-09 | 5.568E-08 | 1.332 (1.146-1.548) | 1.888E-04 | 6.420E-04 |
| T | Snai2 | 2.220E+00 | SNAI2 | 1.238 (1.068-1.434) | 3.969E-03 | 9.997E-03 | 1.285 (1.082-1.527) | 3.589E-03 | 9.387E-03 |
| L | Atp1a3 | 3.272E+00 | ATP1A3 | 1.136 (1.061-1.216) | 6.052E-04 | 1.764E-03 | 1.124 (1.036-1.22) | 8.125E-03 | 1.625E-02 |
| L | Ccl8 | 4.285E+00 | CCL8 | 1.272 (1.142-1.416) | 1.012E-05 | 5.784E-05 | 1.047 (0.926-1.184) | 4.576E-01 | 5.546E-01 |
| L | Cfb | 5.209E+00 | CFB | 1.216 (1.08-1.369) | 6.175E-04 | 1.764E-03 | 1.329 (1.158-1.524) | 2.219E-05 | 1.044E-04 |
| L | Il2rg | 3.515E+00 | IL2RG | 1.207 (1.072-1.36) | 1.531E-03 | 4.083E-03 | 1.185 (1.03-1.364) | 1.565E-02 | 2.846E-02 |
| L | Krt13 | 2.140E+00 | KRT13 | 1.15 (1.085-1.22) | 1.681E-05 | 8.405E-05 | 1.091 (1.007-1.181) | 4.136E-02 | 6.488E-02 |
| L | Krt17 | 3.996E+00 | KRT17 | 1.098 (1.038-1.162) | 1.996E-03 | 4.696E-03 | 1.097 (1.029-1.17) | 7.346E-03 | 1.507E-02 |
| L | Pcbp3 | 2.607E+00 | PCBP3 | 1.17 (1.105-1.239) | 2.537E-07 | 1.908E-06 | 1.14 (1.061-1.224) | 5.423E-04 | 1.669E-03 |
| L | Rac2 | 4.326E+00 | RAC2 | 1.274 (1.109-1.464) | 4.751E-04 | 1.520E-03 | 1.297 (1.099-1.531) | 1.622E-03 | 4.186E-03 |
| L | Saa2 | 3.126E+00 | SAA1 | 1.122 (1.085-1.16) | 5.494E-11 | 2.197E-09 | 1.117 (1.075-1.162) | 6.703E-08 | 7.661E-07 |
| L | Saa2 | 3.126E+00 | SAA2 | 1.118 (1.079-1.159) | 5.374E-09 | 1.433E-07 | 1.105 (1.059-1.152) | 1.184E-05 | 6.315E-05 |
| L | Samsn1 | 2.092E+00 | SAMSN1 | 1.236 (1.074-1.423) | 2.770E-03 | 6.156E-03 | 1.228 (1.039-1.453) | 1.464E-02 | 2.738E-02 |
| L | Serpina3b | 3.226E+00 | SERPINA3 | 1.162 (1.105-1.223) | 9.233E-09 | 1.477E-07 | 1.137 (1.072-1.207) | 3.146E-05 | 1.259E-04 |
| L | Serpina3g | 4.640E+00 | SERPINA3 | 1.162 (1.105-1.223) | 9.233E-09 | 1.477E-07 | 1.137 (1.072-1.207) | 3.146E-05 | 1.259E-04 |
| L | Serpina3i | 5.367E+00 | SERPINA3 | 1.162 (1.105-1.223) | 9.233E-09 | 1.477E-07 | 1.137 (1.072-1.207) | 3.146E-05 | 1.259E-04 |
| L | Slc2a3 | 2.779E+00 | SLC2A3 | 1.217 (1.075-1.379) | 1.756E-03 | 4.256E-03 | 1.257 (1.088-1.452) | 1.676E-03 | 4.191E-03 |
| L | Uck2 | 2.030E+00 | UCK2 | 2.104 (1.705-2.596) | 2.070E-11 | 1.656E-09 | 2.139 (1.664-2.749) | 7.384E-09 | 1.477E-07 |
| L | Wt1 | 3.909E+00 | WT1 | 1.124 (1.064-1.187) | 2.898E-05 | 1.220E-04 | 1.184 (1.109-1.264) | 3.156E-07 | 2.104E-06 |
| KTL | 4930550L24Rik | 4.819E+00 | MAGEC2 | 1.282 (1.182-1.392) | 3.038E-06 | 8.506E-05 | 1.292 (1.146-1.457) | 8.876E-04 | 3.824E-03 |
| KTL | 4930550L24Rik | 4.819E+00 | MAGEC3 | 1.216 (1.119-1.321) | 5.181E-05 | 3.234E-04 | 1.17 (1.05-1.304) | 1.096E-02 | 2.924E-02 |
| KTL | Ccl8 | 3.268E+00 | CCL8 | 1.272 (1.142-1.416) | 1.012E-05 | 1.417E-04 | 1.047 (0.926-1.184) | 4.576E-01 | 5.824E-01 |
| KTL | Cfb | 2.912E+00 | CFB | 1.216 (1.08-1.369) | 6.175E-04 | 3.144E-03 | 1.329 (1.158-1.524) | 2.219E-05 | 2.485E-04 |
| KTL | Dnajc12 | 2.501E+00 | DNAJC12 | 1.246 (1.139-1.363) | 4.973E-06 | 9.282E-05 | 1.24 (1.116-1.377) | 1.441E-04 | 8.069E-04 |
| KTL | Il2rg | 3.381E+00 | IL2RG | 1.207 (1.072-1.36) | 1.531E-03 | 5.360E-03 | 1.185 (1.03-1.364) | 1.565E-02 | 3.984E-02 |
| KTL | Il34 | 3.843E+00 | IL34 | 1.287 (1.137-1.458) | 5.197E-05 | 3.234E-04 | 1.362 (1.176-1.578) | 3.216E-05 | 2.573E-04 |
| KTL | Plac8 | 2.235E+00 | PLAC8 | 1.216 (1.082-1.366) | 1.323E-03 | 4.938E-03 | 1.185 (1.026-1.369) | 2.218E-02 | 5.175E-02 |
| KTL | Serpina3g | 2.509E+00 | SERPINA3 | 1.162 (1.105-1.223) | 9.233E-09 | 5.170E-07 | 1.137 (1.072-1.207) | 3.146E-05 | 2.573E-04 |

**Table S3.**
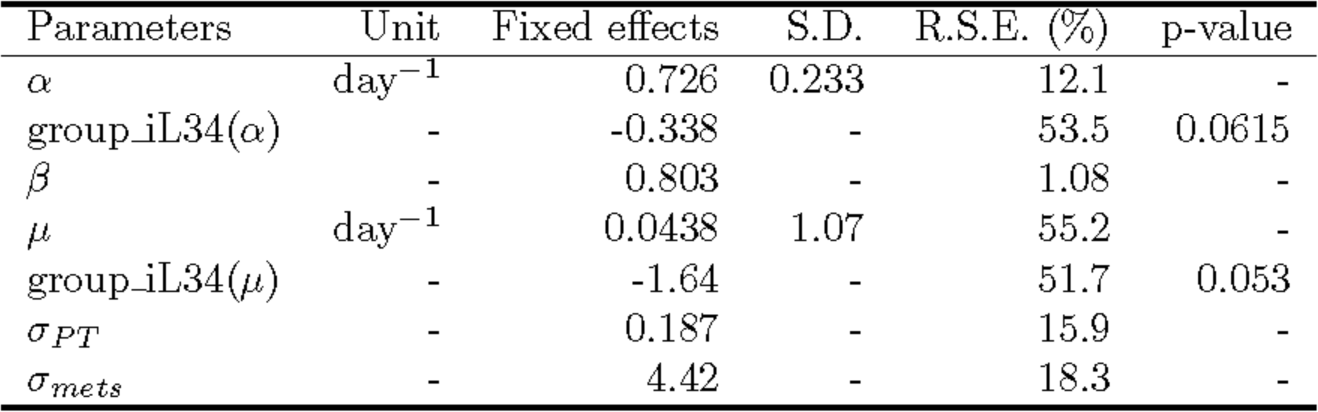
**Parameter values of the mathematical model of primary tumor growth and metastatic dissemination.** A mixed-effects approach was used and associated likelihood maximization for the pooled data (all animals from all groups). Belonging to the IL34 KO group was considered by means of covariates on parameters αand µ. S.D. =standard deviation of the random effects. R.S.E =relative standard error of the estimate. p-value refers to a Wald test for statistical significance of the covariate.

| Parameters | Unit | Fixed effects | S.D. | R.S.E. (%) | p-value |
| --- | --- | --- | --- | --- | --- |
| $\alpha$ | day<sup>-1</sup> | 0.726 | 0.233 | 12.1 | - |
| group_iL34($\alpha$) | - | -0.338 | - | 53.5 | 0.0615 |
| $\beta$ | - | 0.803 | - | 1.08 | - |
| $\mu$ | day<sup>-1</sup> | 0.0438 | 1.07 | 55.2 | - |
| group_iL34($\mu$) | - | -1.64 | - | 51.7 | 0.053 |
| $\sigma_{PT}$ | - | 0.187 | - | 15.9 | - |
| $\sigma_{mets}$ | - | 4.42 | - | 18.3 | - |

## Notes

#### Summary of Updates

We have revised our manuscript to some extent. We made it shorter and more concise. The following modifications have been done: 1/ The abstract was rewritten 2/ We have moved significant parts of the Materials and Methods section to supplementary Materials and Methods 3/ We shortened sections in the manuscript in the Result section and Discussion section to highlight more the important parts of our study and to make the message more transparent. 4/ We removed GRPC5A and PODNL1 from the results and Discussion section because we wanted to emphasize markers in the signatures that were soluble and which could be tested in patient cohorts. 5/ We have corrected some spelling mistakes

